# A Multivariate View of Parallel Evolution

**DOI:** 10.1101/2020.01.26.920439

**Authors:** Stephen P. De Lisle, Daniel I. Bolnick

## Abstract

A growing number of empirical studies have quantified the degree to which evolution is geometrically parallel, by estimating and interpreting pairwise angles between evolutionary change vectors in multiple replicate lineages. Similar comparisons, of distance in trait space, are used to assess the degree of convergence. These approaches amount to element-by-element interpretation of distance matrices, and can fail to capture the true extent of multivariate parallelism when evolution involves multiple traits sampled across multiple lineages. We suggest an alternative set of approaches, co-opted from evolutionary quantitative genetics, involving eigen analysis and comparison of among-lineage covariance matrices. Such approaches not only allow the full extent of multivariate parallelism to be revealed and interpreted, but also allow for the definition of biologically tenable null hypotheses against which empirical patterns can be tested. Reanalysis of a dataset of multivariate evolution across a replicated lake/stream gradient in threespine stickleback reveals that most of the variation in the direction of evolutionary change can be captured in just a few dimensions, indicating a greater extent of parallelism than previously appreciated. We suggest that applying such multivariate approaches may often be necessary to fully understand the extent and form of parallel and convergent evolution.

## Introduction

How repeatable is the evolutionary process? Highpoints in the history of life include taxa that have appeared to independently evolve similar adaptations in response to similar environmental conditions. Such examples represent some of the most striking evidence for adaptive evolution, suggesting not only that evolution can sometimes repeat itself, but that we can also identify the broad environmental factors governing natural selection (Nosil et al. 2002, Langerhans and DeWitt 2004, Losos 2011, Bolnick et al. 2018, Stuart 2019). Thus, this classical question cuts to the core of ongoing debates over the role of chance and determinism in evolution at all timescales.

The question of repeatability in evolution has been reframed in light of contemporary approaches to studying evolution and natural selection in the wild. This new work has distinguished the outcome of evolution, convergent (divergent), from the path of evolutionary change, parallel (nonparallel) (Bolnick et al. 2018). Purpose-built statistical tests have been invented to test hypotheses of parallelism and convergence at both the micro (Collyer and Adams 2007, Adams and Collyer 2009, Collyer et al. 2015) and macro scale (Mahler et al. 2013). This work has led to some important advances in our empirical understanding of how repeatable evolution can be (Oke et al. 2017, Stuart et al. 2017), yet has also highlighted some fundamental challenges (Bolnick et al. 2018) of linking pattern and process in empirical tests of the repeatability of evolution.

Here we emphasize an explicitly-multivariate set of approaches to the study of parallel and convergent evolution. These multivariate approaches surmount three key difficulties: 1) univariate analyses, often employed in studies of parallel and convergent evolution, can be difficult to interpret when the data are multivariate, and can lead to 2) underestimation of the importance of shared dimensions of evolutionary change, which have been difficult to identify due to the challenge of 3) construction of a statistical null expectation that is also biologically appropriate, against which empirical patterns can be pitted. We show how these challenges can be resolved with some basics of linear algebra that permit analysis of entire matrices of similarity of evolutionary change. We illustrate our points by revisiting a published dataset of parallel evolution in a fish species, where a truly-multivariate approach reveals a far greater extent of parallelism than can be concluded from univariate analysis. Our arguments largely ‘parallel’ similar issues raised in a closely aligned subfield, evolutionary quantitative genetics, where it has long been recognized (Lande and Arnold 1983, Phillips and Arnold 1989, Blows and Brooks 2003, Blows 2007a, b, Kirkpatrick 2009, Wyman et al. 2013) that understanding selection on and expression of complex traits requires approaches that are explicitly multivariate.

Below, we first define parallelism and convergence as separate but related patterns. Next, we outline a set of multivariate geometric techniques to explore the degree and form of parallelism, applying these techniques to a reanalysis of a published dataset of parallelism in a fish species. We then review approaches to assessing multivariate convergent/divergent evolution, independent of parallelism, and highlight a specific published case study leveraging such an approach. We discuss advantages and limitations of these multivariate techniques.

## Defining parallelism and convergence as unique phenomenon

Parallel (nonparallel) and convergent (divergent) evolution can be seen as separated but related patterns (Figure 1; see also Bolnick et al. 2018). This separation is potentially important because it is the case that unique evolutionary processes could lead to patterns of evolutionary parallelism without convergence, or vice-versa. But, parallelism and convergence also need not be viewed as mutually exclusive phenomena. For example, conserved directional selection across lineages could result in parallelism without convergence (Figure 1A). Alternatively, adaptation towards a shared optimum by lineages with unique evolutionary histories, and thus unique ancestral positions in trait space, could result in convergence without parallelism (Figure 1B). Yet, parallel evolutionary processes can lead to divergence if, for example, one lineage evolves faster along a shared trajectory. Thus, separating parallelism and convergence may often be necessary to link evolutionary pattern and process.

**Figure 1.**
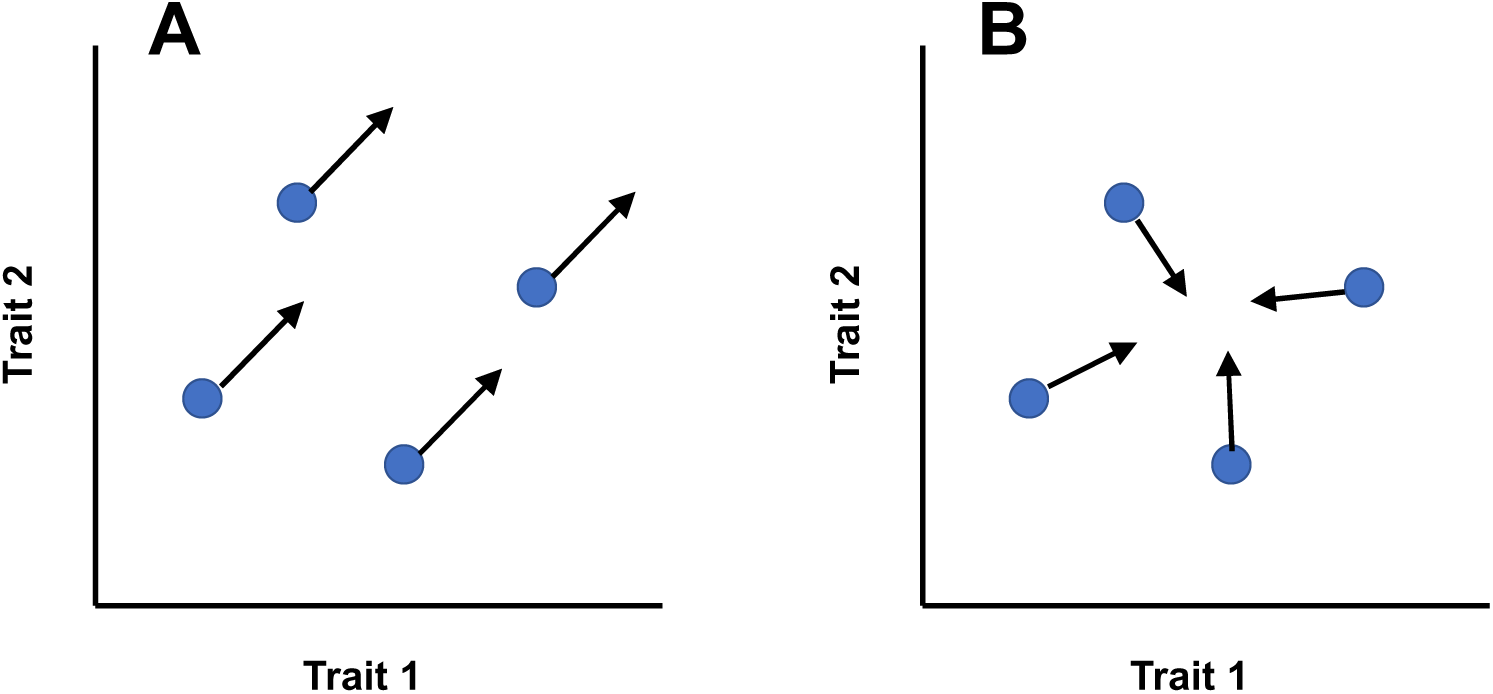
Parallel and convergent evolution as separate but related patterns. Parallel evolution can be defined as a shared orientation in the direction of evolutionary change across two or more lineages, regardless of their starting (ancestral) position, illustrated for four lineages in panel A showing perfect parallelism. Convergent evolution, however, describes a change (reduction for convergence, increase for divergence) of the dispersion of lineages in trait space, illustrated in panel B. Note that evolution can be any combination of parallel or convergent, depending on the ancestral state of the lineages in question and the evolutionary processes at play.

## Geometry of parallel evolution

An intuitive way to define parallelism is via analysis of evolutionary change vectors (Collyer and Adams 2007, Adams and Collyer 2009), where the vector of multivariate evolutionary change across an environmental gradient or time points *a* and *b* (e.g. and ancestral versus descendant population), for a given lineage is

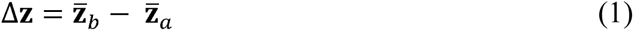

(Lande 1979) perhaps with traits standardized to make units comparable across traits. Note that, beyond morphology, such a vector can be defined using breeding values, sequence variation (Stuart et al. 2017), or gene expression profiles, and that environments can be defined using external abiotic factors or any other feature of interest that defines subpopulations (e.g., sex; De Lisle and Rowe 2017). Δ**z** is a vector of evolutionary change, or evolutionary response to selection, and a growing number of studies have employed an approach (Collyer and Adams 2007, Oke et al. 2017, Stuart et al. 2017) where the angle between Δ**z** vectors is estimated for each pairwise combination of lineages studied, as

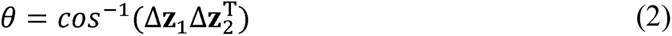

where each vector has been normalized to unit length. Statistical significance of the angle *θ* is often assessed via permutation of the data, comparing the observed value to a null of perfect parallelism across each pairwise combination of lineage replicates (Collyer and Adams 2007, Adams and Collyer 2009). While appropriate for assessing significance of a specific pair of vectors, for example when only two lineages are sampled or are of particular *a priori* interest, this approach is problematic for two reasons when multiple lineages are sampled.

First, how do we interpret values *θ* when more than two lineages are sampled? Lineages would be expected to differ, potentially substantially, in the direction of evolutionary change even when most are evolving in a similar direction. The problem is illustrated in Figure 2A, where three lineages are diverging in multivariate trait space at angles of *θ* = 45.5 degrees from each other. Each pairwise angle interpreted alone suggest evolution is largely nonparallel. Yet this element-by-element interpretation misses the fact that there is a shared common axis of divergence in multivariate trait space that all three lineages load strongly onto and that accounts for a large amount (80%) of the among-lineage variation in the direction of evolution. Moreover, multiple dimensions of parallelism and (anti)parallelism, although a likely feature of multivariate parallel evolution (Figure 2B), cannot be inferred in any intuitive way from interpreting individual angles alone. This challenge in interpreting these pairwise angles individually only becomes greater as the dimensionality of trait space and the number of lineages sampled increases, and is analogous to the within-lineage challenge of using single elements of a genetic covariance matrix to understand the distribution of genetic variation across traits (Kirkpatrick 2009), or the challenge of using single elements of a matrix of nonlinear selection gradients to infer the shape of nonlinear selection (Blows and Brooks 2003, Blows 2007a, b). Put simply, lineages may share a common axis (or axes) of directional change even when most of the individual angular distances between lineages are significantly non-zero.

**Figure 2.**
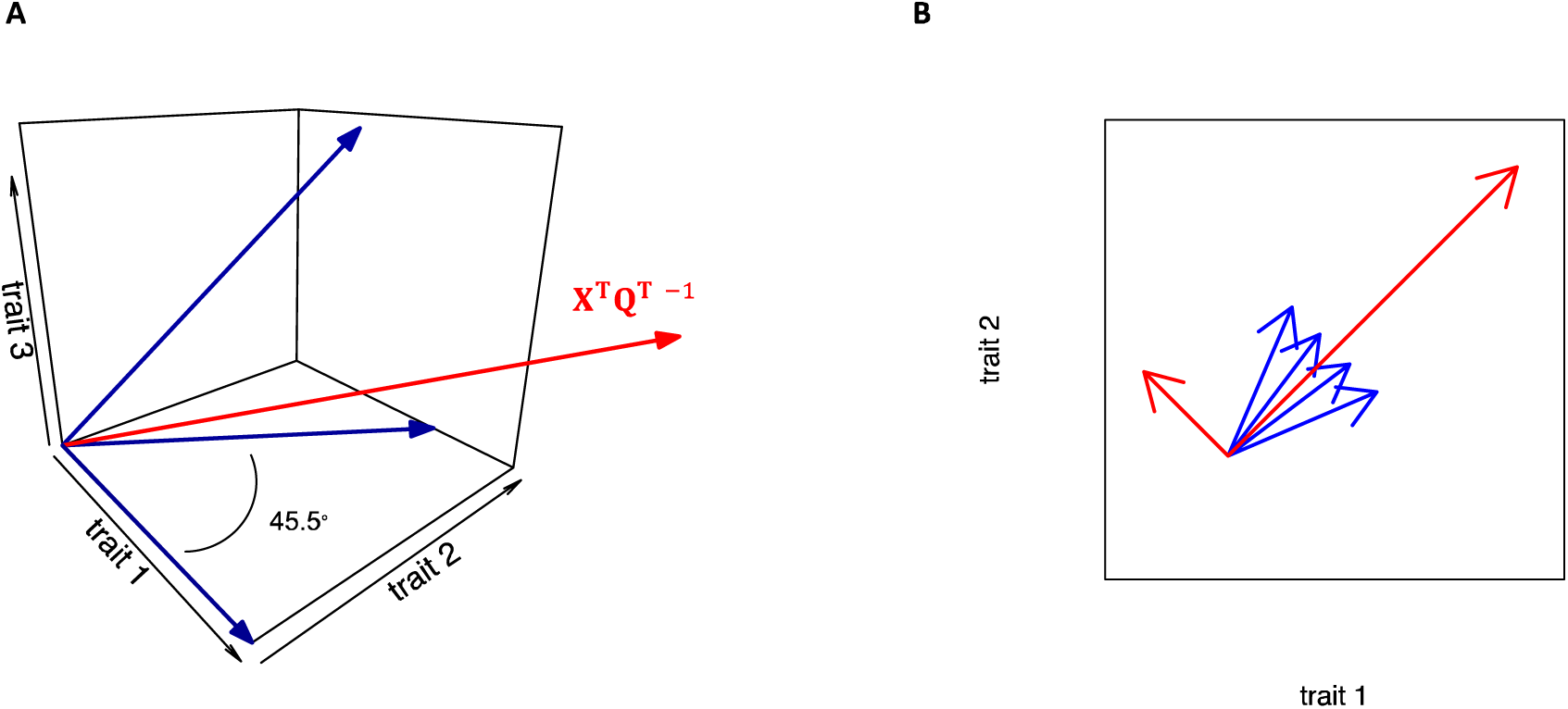
Parallel evolution in multivariate trait space. In Panel A, Vectors of evolutionary change for three traits sampled from 3 lineages are plotted as blue arrows. Each pairwise combination of lineage vectors differ in orientation by 45.5 degrees in three dimensional trait space. Analysis of these angles alone would indicate evolution is only weakly, if at all, parallel. Yet all three lineages load strongly on a single major axis of evolutionary change that accounts for 80% of the among-lineage variation in the orientation of evolution. The red arrow plots this vector of shared multivariate parallel evolution back into three dimensional trait space (see eqn 3-5). Panel B illustrates a similar scenario but sampling four lineages diverging in two-dimensional trait space. Red arrows plot the two orthogonal vectors of shared multivariate parallel evolution, indicating one major axis of parallel evolution upon which all four lineages load positively, in addition to a second axis upon which some lineage evolutionary change vectors load positively, and some negatively, indicating antiparallel evolution in this direction across the sample of lineages.

A second and related difficulty with interpreting angular distances individually is the problem of what the null is for the angle between any given pair of lineages (Bolnick et al. 2018). Many investigators may be interested in comparing observed patterns to a null of independent directions of evolution across lineages. This null results in an expected value of 90 degrees for any pair of lineages. Yet intuitively, the null *distribution* should depend on both the number of traits sampled and the total number of lineages (i.e., beyond any given pair in question), as sampling many lineages evolving independently in low-dimensional trait space is expected to often result in some highly parallel pair-wise combinations through chance alone. Moreover, investigators may be interested in testing more sophisticated null hypotheses, for example comparing patterns of parallelism to what may be expected due to biases from genetic correlations when estimate(s) of **G** are available (Schluter 1996), and it will often be difficult to define such a corresponding null angle that can be tested using the permutation approach typically used in studies of parallelism.

These challenges emerge from what is, at its essence, an element-by-element approach to analysis of angular distance matrices, and they can be circumvented by adopting approaches that consider explicitly the entire matrix of similarity among lineage’s evolutionary change vectors. We can define a matrix of Δ**z** of vectors for *n* traits from *m* lineages as

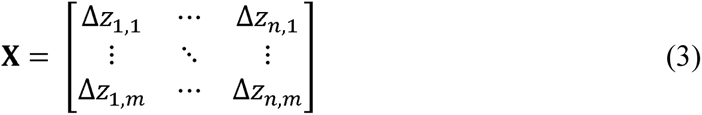

with *m* rows as replicate lineages and *n* columns as traits. From this data matrix, where each element is the evolutionary change value for a single trait from a single independently evolving lineage, and each row thus represents a Δ**z** vector from a given lineage, we can calculate two potentially relevant covariance matrices. One is the *n*×*n* matrix (where *n* is the number of measured traits) of among-lineage variances and covariances in the traits’ evolutionary changes

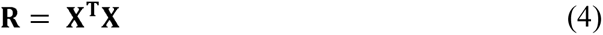

This captures the variances and covariances of evolutionary change across replicate lineages for each trait. Spectral decomposition (eigen analysis) of **R** would be relevant for understanding parallel evolution, since parallel evolution would lead to dimensions of **R** with very low variance. However, it is a challenge to test hypotheses about null or nearly-null dimensions (Kirkpatrick 2009, Blows and McGuigan 2015). So, for the specific question of, “*How parallel is evolutionary change?”*, we may be better off focusing on a different product of **X**,

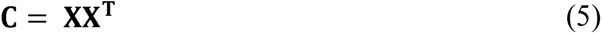

Which is the matrix of vector correlations (assuming the rows of **X** have been normalized) of evolutionary change vectors for each pair of independent lineages in **X**. Thus **C** is a *m*×*m* correlation matrix where *m* is the number of replicate lineages. This matrix contains ones on the diagonal, and each off-diagonal describes the correlation between multivariate evolutionary change vectors across a pair of lineages. By describing the relationship between lineage vectors as a correlation instead of an angle (i.e., a distance), we can use the spectral decomposition (eigen decomposition) of **C**,

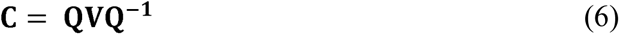

where **Q** is the matrix of eigenvectors and **V** is the diagonal matrix of eigenvalues, to assess two major questions on the degree and form of multivariate parallelism that cannot be tested using univariate approaches:

1. *How extensive is parallelism? Parallel evolution (and anti-parallel) will be captured by the leading eigenvector(s) of C.* In principle at the extreme, where all evolutionary change is perfectly parallel, **C** would be of unit rank, with all variance (= *m*) captured in a single dimension, ***q_1_***, of shared evolutionary change. At the other extreme, if evolutionary change is completely independent in directional across all lineages sampled, **C** would be of rank equal to *m* or *n* (whichever is lower), with variance distributed uniformly across non-null (if more lineages than traits are sampled) eigenvectors. Thus, the strength of parallelism captured by the leading eigenvector (***q_1_***) of **C** can be expressed as *ν*_1_/∑*ν*, where *ν*_1_ is the leading eigenvalue of **C.** True parallelism is reflected when all lineages load positively on the corresponding eigenvector ***q_1_***, while anti-parallelism would be reflected by a mixture of positive and negative loadings on ***q_1_***.
2. *How many dimensions of shared evolutionary change exist? (Anti)Parallel evolution in multiple orthogonal directions would be reflected in multiple significant eigenvalues.* When multiple dimensions are identified as significant (see below), these dimensions reflect orthogonal axes of parallel and anti-parallel evolution. Multiple dimensions suggest more than one alternative solution to a particular adaptive challenge, or evolution towards more than one phenotypic optimum, and hence reflect parallel evolution among certain comparisons between replicate descendant populations but non-parallel evolution for other such comparisons. Figure 2B illustrates such a scenario.

Thus, the eigenvectors and eigenvalues of **C** provide information on both the extent of multivariate parallelism and contributions of specific lineages to parallel evolution or lack thereof. Eigenvector scores could, in principle, be related to other variables of interest, such as among-lineage environmental variation via canonical correlation or other approaches. We can also relate the eigenvectors of **C** back to trait space via

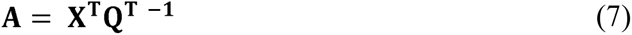

Where **A** is a matrix with *m* column vectors of length *n* relating each eigenvector of **C** back to trait space (Figure 2). We can then see how traits load on these vectors to establish which traits are contributing most to parallelism.

Of course, sampling error will create variance in all or most dimensions. This same sampling error will also lead to a skewed distribution of eigenvalues even in the absence of actual parallel evolution (Johnstone 2001). This leads to the problem of constructing a null expectation for evolutionary parallelism against which empirical patterns can be tested. Specifically, what is the null expectation for the distribution of eigenvalues of **C**? This is a problem in random matrix theory, where the distribution of eigenvalues of random covariance matrices has been of great interest (Tracy and Widom 1996, Johnstone 2001, Tracy and Widom 2009, Blows and McGuigan 2015, Sztepanacz and Blows 2017). Generally, the leading eigenvalue of a sample covariance matrix is expected to be follow a Tracy-Widom distribution (Tracy and Widom 1996, 2009, Blows and McGuigan 2015, Sztepanacz and Blows 2017); however, this distribution is sensitive to centering and scaling constants that may make hypothesis testing difficult (Blows et al. 2015), and we may also be interested in assessing significance of more than one dimension (as suggested above). A null distribution of all *m* eigenvalues can be obtained by sampling the *m*-dimensional Wishart distribution with *n* df, where the hypothesis being tested depends on the covariance structure of the sampled Wishart distribution (Johnstone 2001). If we are interested in testing a null hypothesis of independent evolution across lineages, this would correspond to zero off-diagonals (an identity matrix) in the Wishart covariance structure. An example of such a null distribution under varying *n* is illustrated in Figure 3. For the case of more lineages than traits, we can simulate random vectors to establish the null empirically, although the strongly skewed distribution of eigenvalues in such a scenario indicates that exceptionally strong parallelism is required to reject the null (Figure 3). By comparing the eigenvalues of **C** to the expected distribution of eigenvalues under the null expectation, we can establish whether evolution is significantly more parallel in multivariate trait space, than null expectations. And, we can test whether this is true in more than one dimension, which would be reflected in multiple eigenvalues greater than the null expectation.

**Figure 3.**
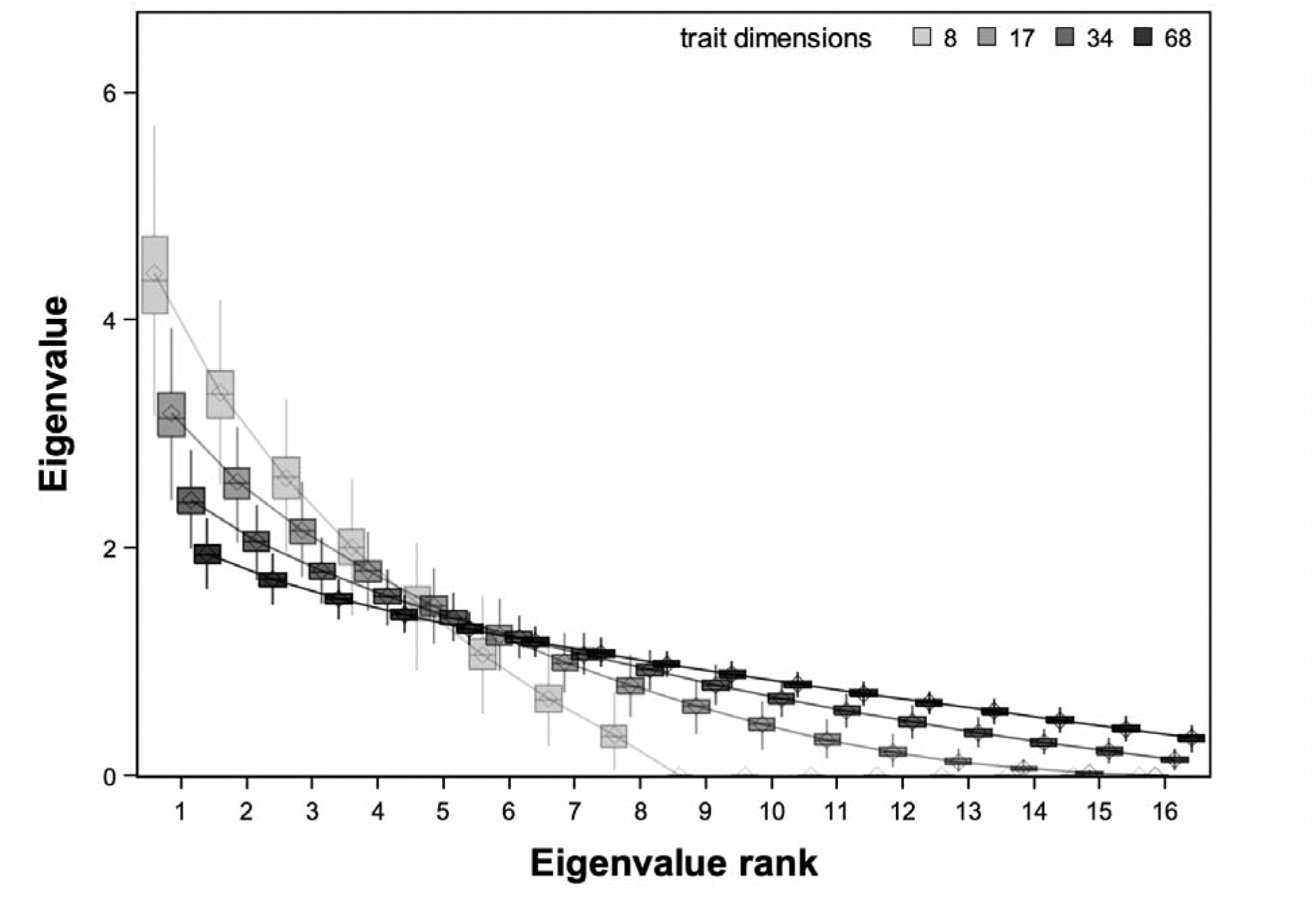
Null distributions of eigenvalues for a 16-dimensional (16 lineage) C matrix. Increasing trait dimensionality relative to lineage replication leads to null distributions favorable to uncovering parallelism. Under the null hypothesis of no relationship between vectors of evolutionary change, which would be reflected in a perfectly uniform distribution of eigenvalues, correlations between lineages (manifest as eigenvalues exceeding unity) are still expected to be observed through sampling error. The magnitude of these expected leading eigenvalues increases with decreasing trait dimensionality simply due to random sampling. For example, when the number of traits is less than the number of lineages (vectors) sampled, strong correlations are inevitable even when the null hypothesis of random evolutionary divergence across lineages is true. Thus, sampling many traits relative to lineages flattens the null distribution of eigenvalues, increasing power to distinguish true parallel evolution from what is expected purely from sampling error under the null hypothesis. It is important to note, however, that high trait dimensionality simply creates null distributions favorable to uncovering parallelism; choice of traits to include in any analysis is primarily a biological problem. Distributions obtained by sampling the corresponding Wishart distribution, with the exception of 8 traits which was constructed empirically by placing random vectors in trait space. Dimensionality chosen here for continuity with the empirical example from Stuart et al. 2017.

Although a null hypothesis of independent directions of evolution among lineages is likely to be the most appropriate null for many tests of the presence and importance of parallelism, other null hypotheses may be of interest. For example, one may wish to test empirical patterns against a null of perfect parallelism, when researchers seek to identify population-specific (e.g., non-parallel) aspects of evolution. In such cases a null distribution of unit rank matrices would be appropriate. Alternatively, we may also be interested in the degree to which genetic covariances and selection shape parallelism. We can relate **C** to **G**, for the case of *m* < *n*, by postmultiplying the right hand side of equation 5 by the *m* dimensional identity matrix in the form of 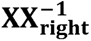 and realizing that under directional selection and constant **G**, **R ≈ GBG** (Zeng 1988), where **B** is the covariance matrix of directional selection gradients across lineages. However, because both **G** and **R** share the same dimensionality (number of traits, as opposed to lineages as with **C**) it may be far more straightforward to test hypotheses on the influence of **G** on evolutionary change via analysis of **R** instead of **C** (e.g., see Chenoweth et al. 2010). It is likely that constructing more sophisticated null expectations for contrasts in orientation may require development of theoretical models that focus specifically on the geometry of divergence in complex traits (e.g. see Thompson et al. 2019, whose model is constructed specifically to bear relevance to angular contrasts).

## Example: Parallel evolution across a lake-stream gradient in threespine stickleback

Threespine stickleback in postglacial inland waters of the Northern hemisphere represent one of the most compelling models for testing the repeatability of evolution. Repeated colonization of freshwater from marine environments has resulted in replicated divergence from a marine ancestral phenotype into multiple freshwater forms (McPhail 1993, Bell and Foster 1994, Colosimo et al. 2005). In a comprehensive study of parallel evolution in this system, Stuart et al. (2017) sampled morphology, genetics, and environmental variables from lake and stream subpopulations from 16 lineages (watersheds) on Vancouver Island, Canada. They found only limited evidence of parallelism in morphological divergence in 84-dimensional trait space between lake and stream subpopulations across the 16 replicate lineages, as inferred in part from a relatively even distribution of angular distances across all 120 pairwise lineage combinations (their figure 3b) with no clear signal of a skew towards values near zero. By correlating these pairwise angular distances in morphological change vectors, with estimates of angular distance in genetic and environmental change vectors, Stuart et al. showed that deviations from parallelism among lineages is due to non-parallelism in environmental factors, and levels of gene flow across subpopulations. This study is unique in leveraging additional data to explain variation in morphological parallelism, yet the high dimensional nature of the Stuart et al. data exemplifies the challenges of inferring the overall degree of parallelism from angular distance alone.

Although element-by-element interpretation of angular distances provides little evidence of strong morphological parallelism in the Stuart et al. data, there are a few traits that were strongly parallel. In a reanalysis of their morphological dataset, we find that spectral decomposition of the matrix of vector correlations in morphological change vectors, **C**, reveals a strongly skewed distribution of eigenvalues (Figure 4). Comparison of this distribution to that expected under a null hypothesis of random (in direction) evolutionary change across lineages reveals strong statistical support for three dimensions of parallel evolution (Figure 4). The leading eigenvalue is greater than 7, capturing nearly 50% of the variance among lineages in the direction of evolution, and together the three significant dimensions of parallelism explain 77% of the among-lineage variation. Examination of the lineage loadings on the corresponding eigenvectors reveals that the sign of these loadings is shared by most lineages, indicating these shared axes of divergence are largely (although not completely) axes of parallelism, as opposed to antiparallelism (Figure 5). Projecting these vectors back into trait space (via equation 7) indicates that traits classified as those related to defense, swimming, and trophic interactions load most strongly on these three axes, compared to traits that are generally unclassified (Figure 6). This analysis (SAS/IML script as well as estimates of **C**, **Q**, **V**, and **A**, provided in the supplemental material) suggests an important role for morphological parallelism across lineages, and suggests traits known to play a role in performance, defense, and resource acquisition to be important in parallel adaptation across a lake/stream gradient.

**Figure 4.**
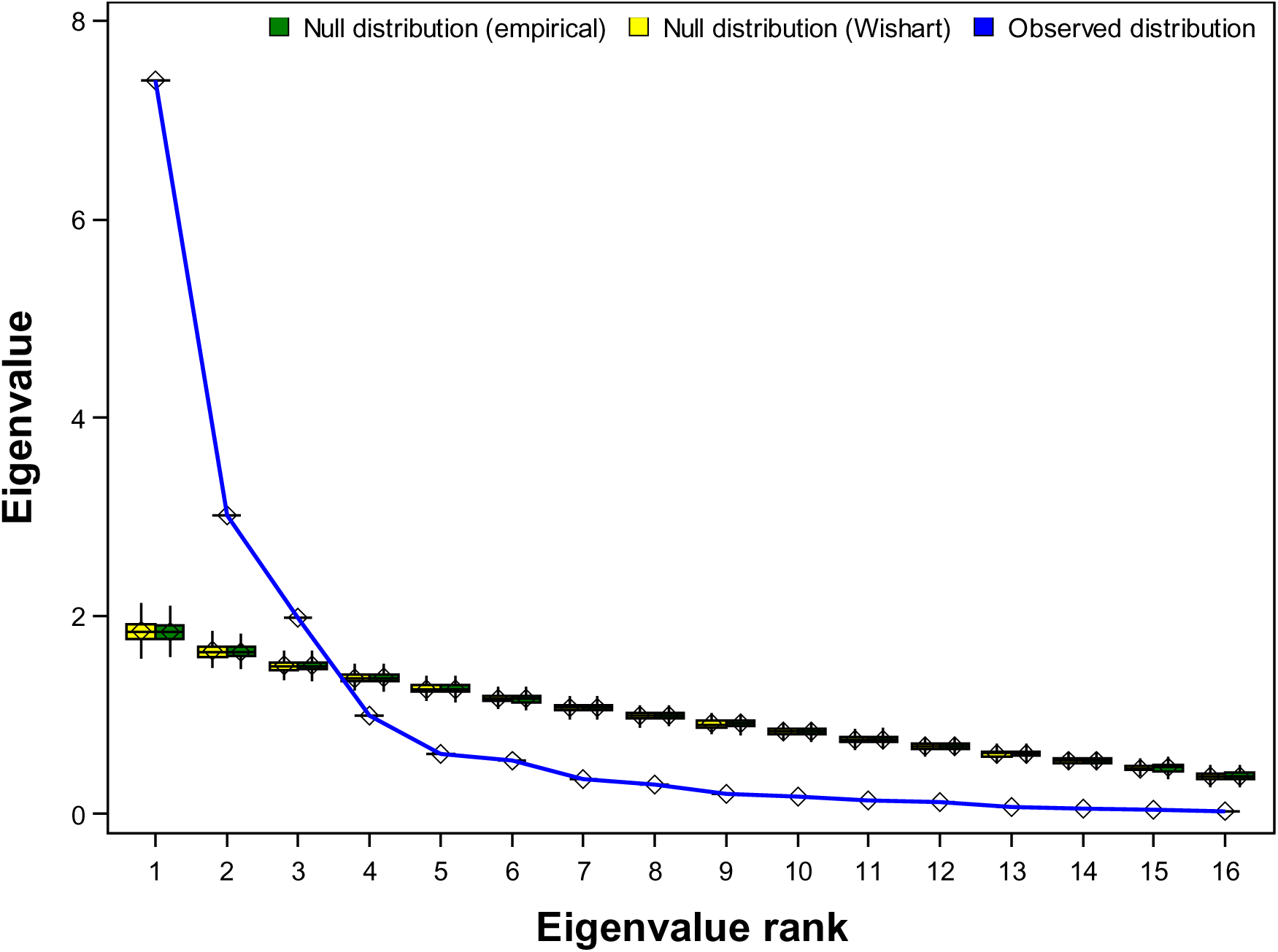
Distribution of eigenvalues from matrix C from Stuart et al. 2017 data, showing statistical support for three orthogonal dimensions of parallel change. Yellow and green boxplots indicate the expected distribution of eigenvalues under a null hypothesis of a random direction of evolutionary change cross lineages (zero off-diagonals in the matrix of vector correlations), calculated from sampling the corresponding Wishart distribution or empirically by placing random vectors in trait space (each 1,000 X). Blue indicates values observed from diagonalization of the estimated matrix of vector correlations, **C**. **C** was calculated from Supplementary data Table 1 in Stuart et al. (2017), with each row of **X** first normalized to unit length.

**Fig 5.**
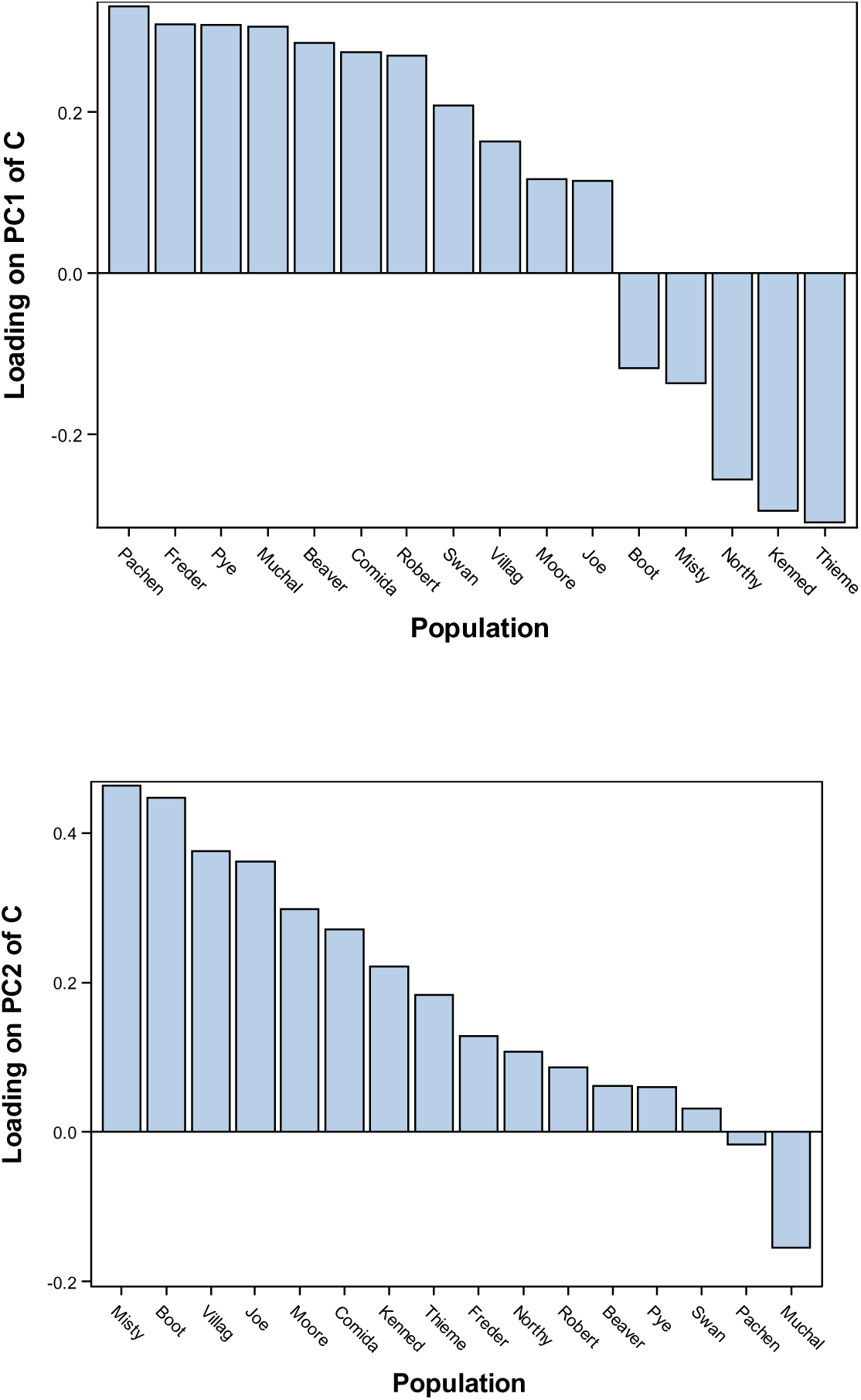
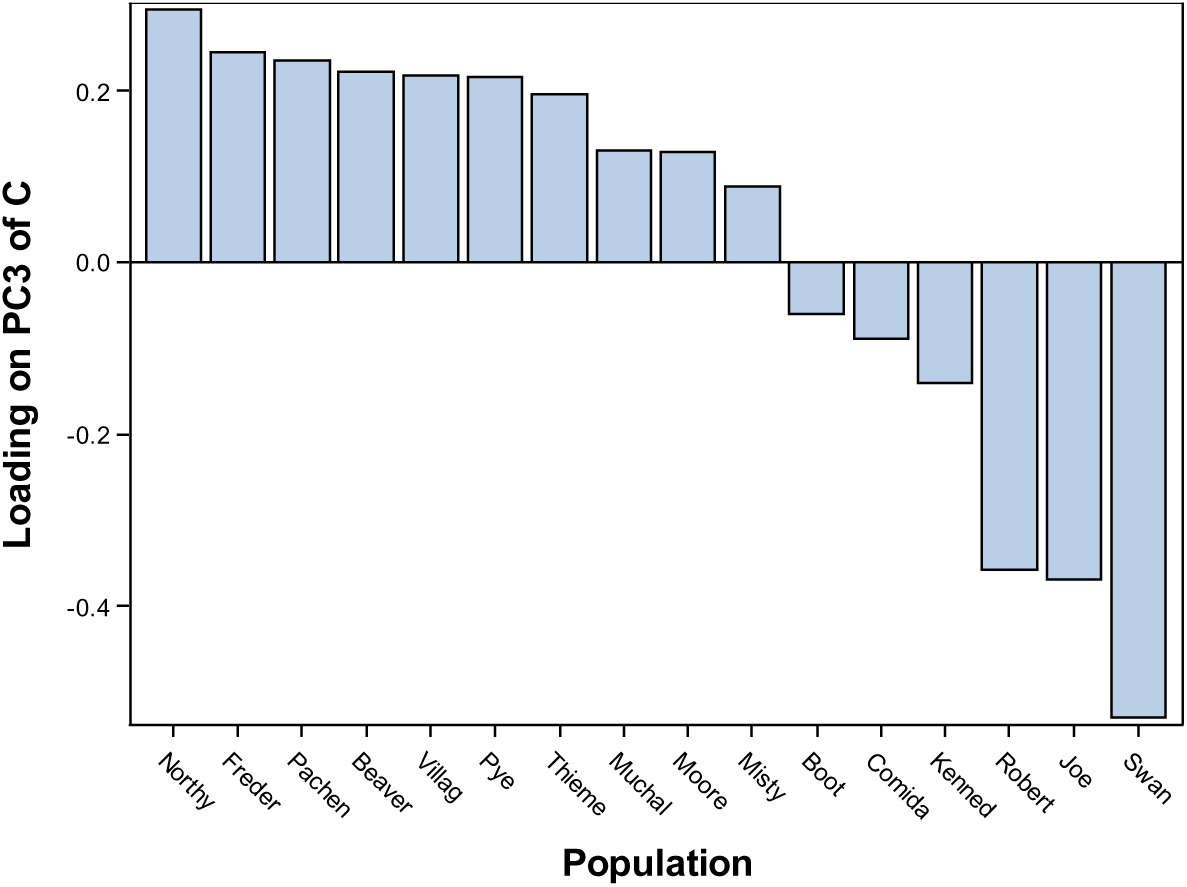
Loadings for each lineage on the first three eigenvectors (principle components) of C. The observation that most lineages are loading positively on PC1 and PC2 indicate that those are dimensions of (mostly) true parallel evolution, as opposed to anti parallel evolution, which would be represented as a mixture of positive and negative loadings across lineages, for example PC3.

**Fig 6.**
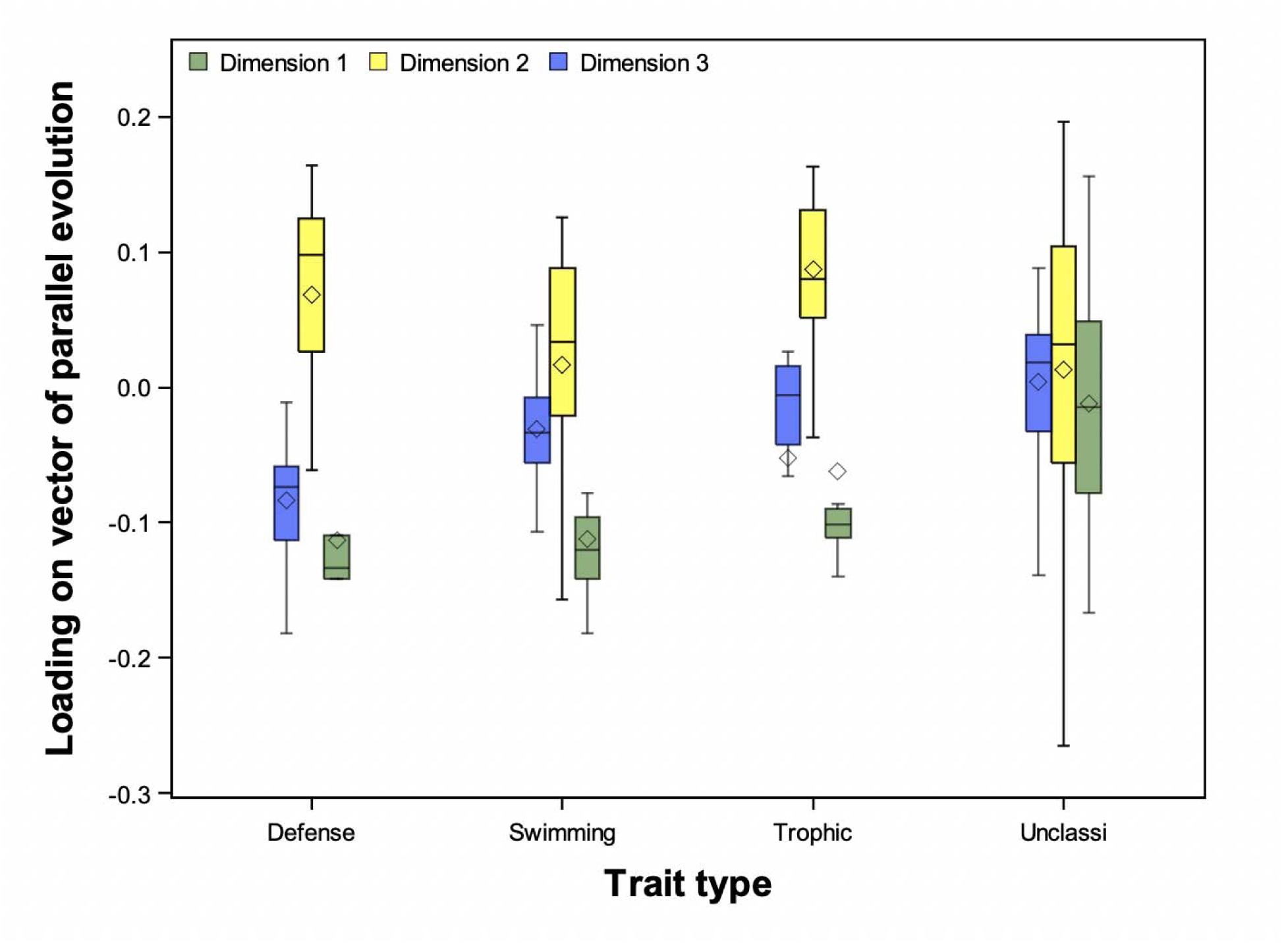
Each of the three significant dimensions of parallelism, estimated from reanalysis of Stuart et al. 2017, related back to trait space. Trait classifications are taken from Stuart et al. 2017.

By revealing statistical support for three dimensions of (imperfect) parallelism, our revisit of the Stuart et al. dataset in some ways recapitulates their findings: we do not find evidence of a single dimension of complete parallelism, which would be reflected in only a single significant eigenvalue with an eigenvector that all lineages load positively onto. Thus, patterns of evolutionary change in this system are apparently complex, and influenced by more than the simple environmental classification of ‘lake’ versus ‘stream’, and the analyses in Stuart et al. (2017) show how such deviations from parallelism can be associated with potential causal sources of variation. However, the multivariate approach we take here suggests a far more important role for morphological parallelism than can be inferred from interpretation of individual angular distances: most of the variation in evolutionary change in 84 dimensional trait space across 16 independent lineages can be captured in just three dimensions, indicating overwhelming statistical support (c.f. lead eigenvalue in Figure 5 to null) to reject a null hypothesis that evolution has proceeded in random directions in trait space across these lineages. Moreover, although trait-by-trait univariate linear models suggest that the traits with clearly-defined ecological functions rarely diverge in parallel in univariate space (Stuart et al.’s Figure 1A), our multivariate analysis reveals that these traits are in fact the most important contributors to shared axes of parallel divergence in multivariate trait space (Figure 6). This is consistent with biological intuition that traits related to swimming performance, defense from predators, and feeding may play an important role in adaptation to lake versus stream environments. Thus, our reanalysis of the Stuart et al. dataset illustrates the utility of taking a multivariate approach to assessing the overall degree of parallelism, an approach which complements their original analyses relating deviations from perfect parallelism to gene flow and environmental variation.

## Convergent and divergent evolution in multivariate trait space

We can think of convergent versus divergent evolution as a distinct and non-exclusive phenomenon from parallelism (Figure 1; Bolnick et al. 2018). Past workers have proposed separate (from analysis of parallelism) tests of convergence via estimation of the distribution of pair-wise differences in the lengths of evolutionary change vectors

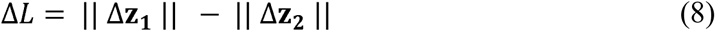

calculated for each pairwise combination of lineages, where non-zero values imply some degree of convergence or divergence (Collyer and Adams 2007, Stuart et al. 2017, Bolnick et al. 2018). Alternatively, estimation of the change in Euclidean distance between lineage pairs across environments/timepoints *a* and *b*

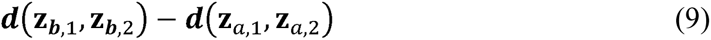

can be calculated, where negative values would indicate convergence between a given lineage pair and positive values divergence (Bolnick et al. 2018), and this approach could perhaps be modified by instead calculating Mahalanobis distance standardized by a pooled covariance matrix. The former approach (equation 8) is problematic because it is difficult to interpret how values of Δ*L* correspond to convergence vs divergence (Bolnick et al. 2018). The latter approach is essentially an indirect comparison of the variance among lineages in one environment versus the other, which captures multivariate convergence versus divergence in an intuitive, albeit somewhat indirect (when more than two lineages are sampled) way. Both approaches are problematic in that they do not allow assessment of which traits contribute most to convergence versus divergence, nor do they account for more complex scenarios where the form of convergent/divergent evolution differs across traits or trait combinations (Figure 7).

**Figure 7.**
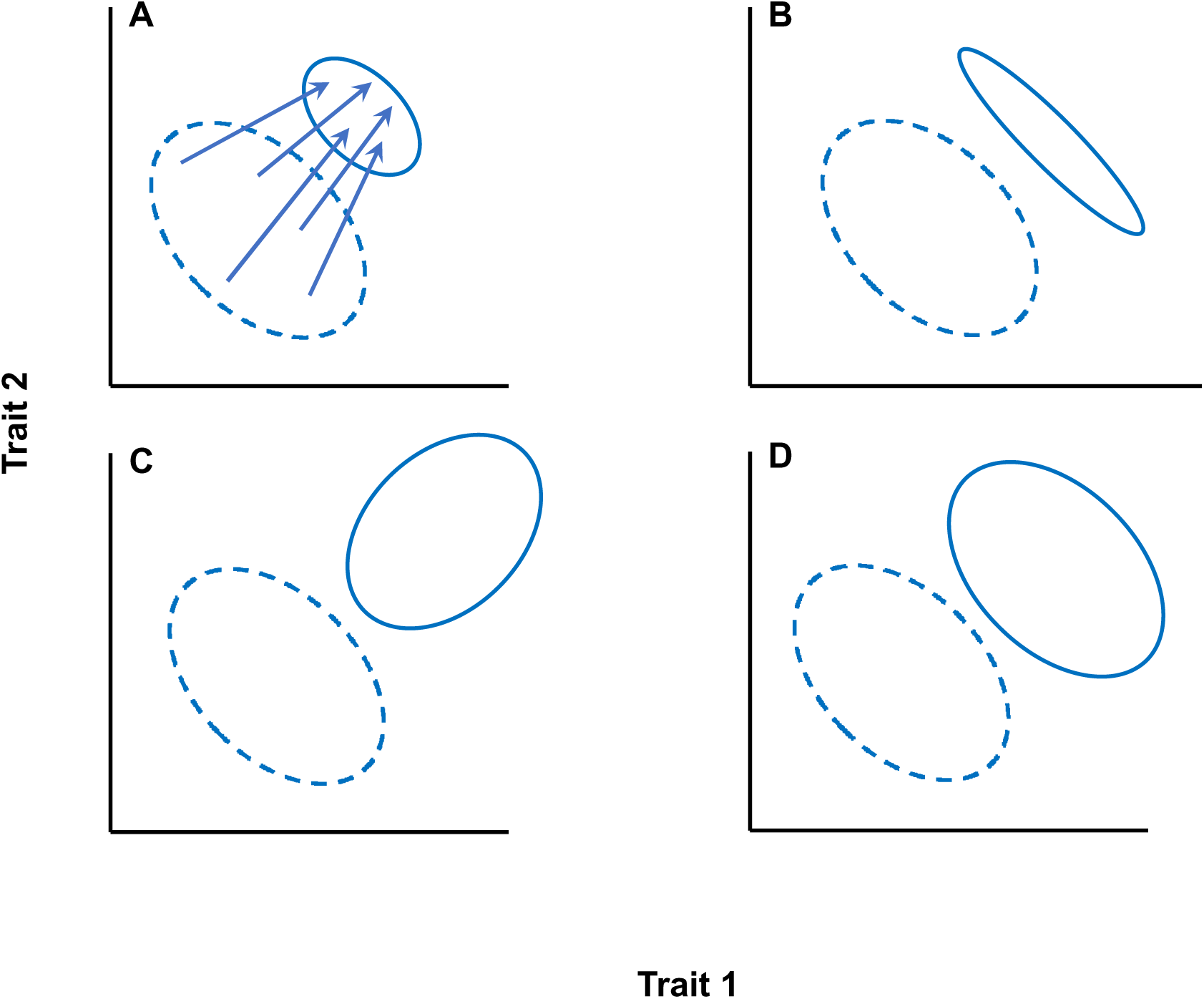
Convergent and divergent evolution in multivariate trait space. Comparison of among lineage variance-covaraince matrices **D** of mean trait values estimated at timepoint (or environment) one (dashed elipses) and two (solid elispses) indicate the form of convergent or divergent evolution. Arrows in panel A illustriate hypothetical evolutionary change vectors, which are left off of subsequent panels for concision. In panel A, evolution is net-convergent and all traits contribute to this convergence; matrices are proportional. In panel B, evolution is convergent in only a single trait dimension. In panel C, evolution is both divergent and convergent, in different combinations of traits, leading to changes in the orientation of the among-lineage covariance matrix. In panel D, evolution is neither convergent nor divergent in any combination of traits, despite strong parallel change across all lineages, leading to no net change in the among-lineage covariance. All cases illustrate largely parallel evolutionary change in overall size.

A more direct approach is to compare the among-lineage covariance matrices of trait mean values, **D** (Lande 1979, Lande 1980), estimated separately for the two environments or timepoints. Given we have vectors of trait means **z***_a_* and **z***_b_* corresponding to timepoint/environment a and b for each lineage, we can calculate their separate among lineage covariance matrices **D***_a_* and **D***_b_*. Multivariate convergent evolution would result in a reduction in among-lineage variance, and a reduction in the size of the covariance matrix **D**. Thus a comparison of the traces of **D***_a_* and **D***_b_* provides a test of whether evolution has been net-convergent or net-divergent, similar to the analysis of pairwise Euclidean distances as in equation 10.

The comparison of the among-lineage covariance matrices provides potential for additional insight when patterns of selection or genetics produce convergence in some combinations of traits, and divergence in others (Figure 7). Although the formal comparison of covariance matrices is a complex statistical problem, the challenge is reduced when only a single pair of matrices is to be compared (as is the case when only two environments or time points are studied) and when one is interested in retaining and contrasting all principle components (Blows et al. 2004). In this case, a Common Principle Component Analysis (CPCA) approach allows for a straightforward test of how **D***_a_* and **D***_b_* differ that is also easily interpretable (Flury 1988, Phillips and Arnold 1999). In the simplest case, a nested multivariate mixed effects model can be used to test the null of **D***_a_* = **D***_b_*, rejection of which indicates convergence (or divergence) in some combination of traits.

Finally, we note that trait scaling represents a particular challenge in analysis of convergence regardless of analytic approach. What do we do when traits are expressed in different units, or differ substantially in value, or evolvability? In some cases, analysis of the raw trait values (population means) may appropriate. Alternatively, one could center the population means and scale them by a pooled estimate of evolvability, such as the pooled within-population phenotypic or genetic covariance matrix **P** or **G**, if available. Regardless, we emphasize that the issue of trait scale is a complex and critical topic (see Houle et al. 2011), that should be carefully considered in studies of multivariate evolutionary change.

## Example: Multivariate divergent evolution during adaptation to a novel environment

Here we provide no novel analyses, but rather highlight a published study employing the approaches outlined above to test hypotheses related to multivariate convergent evolution. Although a number of studies have analyzed eigenstructure of **D** matrices to test hypotheses on the form of divergence across lineages (Schluter 1996, Blows and Higgie 2003, McGuigan et al. 2005, Hohenlohe and Arnold 2008, Kolbe et al. 2011, Punzalan and Rowe 2016, De Lisle and Rowe 2017), the study of Schoustra et al. (2012) provides an example of how formal comparison of **D** matrices across environments provides a straightforward approach to testing hypotheses related to multivariate convergence. In this study the authors used a laboratory experimental evolution approach to understand how population size influences convergent/divergent evolution. To do so they adapted replicate lineages of filamentous fungus (*Aspergillus nidulans*), originating from a common ancestor, to a novel laboratory environment under two population size treatments, low and high. They then measured four traits related to colony formation at the conclusion of 800 generations of adaptation. They used a nested multivariate mixed effects model and found no evidence to reject the null of **D***_low_* = **D***_high_*, indicating no evidence of multivariate convergence/divergence across population size treatments. They confirmed this result using CPCA, finding no evidence to reject a null of matrix equality in favor of proportionality. Finally, Schoustra et al. (2012) used a factor analytic mixed modelling approach to estimate the rank of the pooled **D** matrix, finding statistical support for a full rank divergence matrix. Together, these analyses indicate divergent evolution in all trait dimensions following adaptation from a common ancestor, and that this divergence was not influenced by population size environments. Importantly, this study illustrates how analysis of **D** matrices in a study of convergent evolution allows for biological hypotheses related to multivariate convergence to be defined in terms of specific statistical contrasts of **D** across environments of interest.

## Discussion and Conclusions

Here we advocate for an explicit multivariate approach in studies of parallel and convergent evolution. Spectral decomposition and comparisons of two categories of covariance matrices, the matrix of vector correlations between lineage evolutionary change vectors (**C**) and the among-lineage covariance matrices in trait mean values (**D**), allow biologists to test hypotheses concerning the extent and form of multivariate parallel or convergent evolution (respectively), and to identify the crucial actual lineages and traits underlying these patterns. Whenever multiple traits from multiple lineages are sampled, such an approach provides a potentially more complete picture of the degree and extent of evolutionary change than does element-by-element analysis (e.g., pairwise angles), as is often employed in such studies.

The general approaches we advocate are certainly not new to evolutionary biology, having been developed, described, and applied extensively in the evolutionary quantitative genetics literature. Moreover, such approaches are likely already conceptually grasped by workers familiar with the mechanics of a principle component analysis, and are readily implemented in existing software capable of performing simple matrix manipulations (e.g., R, SAS/IML, Matlab) and perhaps fitting of mixed effects models. Relevant purpose-built statistical packages are also available (e.g. Melo et al. 2015). The point of our paper, then, is simply to highlight one growing (Bolnick et al. 2018, Stuart 2019) subfield of evolutionary biology – quantitative studies of evolutionary parallelism – where these multivariate approaches may be particularly useful but are often not yet employed. Indeed, in many cases ascertaining the true degree and form of parallel evolution, in particular, may be otherwise impossible.

We have suggested that spectral decomposition of among-lineage covariance matrices may often indicate a greater extent of parallelism than is implied through element-by-element interpretation of angular distances. Our reanalysis of a high dimensional dataset of morphological evolution in stickleback lineages support this suggestion, and indicate a greater role for parallelism than can be concluded from individual interpretation of angular distances (Stuart et al. 2017). Of course, whether this suggestion is broadly true is an open empirical question requiring additional studies from a variety of taxa. Moreover, our work suggests that in some cases, especially when trait dimensionality is low, comparison of the eigenvalues of **C** to an appropriate null may indicate little support for parallelism where an element-by-element approach would suggest otherwise. This is because large correlations between evolutionary change vectors from independent, randomly-diverging lineages are expected when many lineages are sampled relative to the number of traits measured. Although high trait dimensionality often presents a challenge in analysis of multivariate datasets, analysis of vector correlations transposes the dataframe (*cf.* equations 4 and 5) in such a way that flips the burden of statistical power to favor sampling of traits. However, an important caveat is that any test of convergent versus divergent evolution via comparison of **D** matrices, as also suggested here and illustrated in past studies (e.g. Schoustra et al. 2012), will favor sampling of lineages over traits. This suggests that careful consideration of the question of interest (tests of parallelism or tests of convergence) may be critical in the early stages of study design if the approaches advocated here are to be employed. Alternatively, in some cases it could be desirable to identify the traits contributing most to parallelism via eigen analysis of **C**, and then focus on contrasts of lower dimensional **D** matrices estimated using only a subset of key traits. Ultimately, the decision of number and type of traits included in any analysis is a biological, rather than statistical or geometric, problem; although high trait dimensionality results in null expectations favorable to uncovering parallelism, including traits arbitrarily may weaken the effects observed through parallel divergence in the traits that matter. Conversely, choosing traits non-randomly can amount to cherry-picking that can overemphasize parallelism.

Analysis of the orientation of evolutionary change, either through angular distance or vector correlation, has intuitive appeal for the quantification of parallelism and repeatability in evolution. When evolutionary change involves multiple traits sampled across multiple lineages, the question of *how repeatable is the evolutionary process?* is one that will often transcend both individual traits and individual lineages, and so will be most clearly addressed with the multivariate approaches presented here.

## Supporting information

Supplemental SAS Script

## Acknowledgements

We are grateful to Yoel Stuart and David Punzalan for discussion and their helpful comments on the manuscript. Funding was provided by the University of Connecticut to DIB.

## Supplemental Material

**Table S1.**
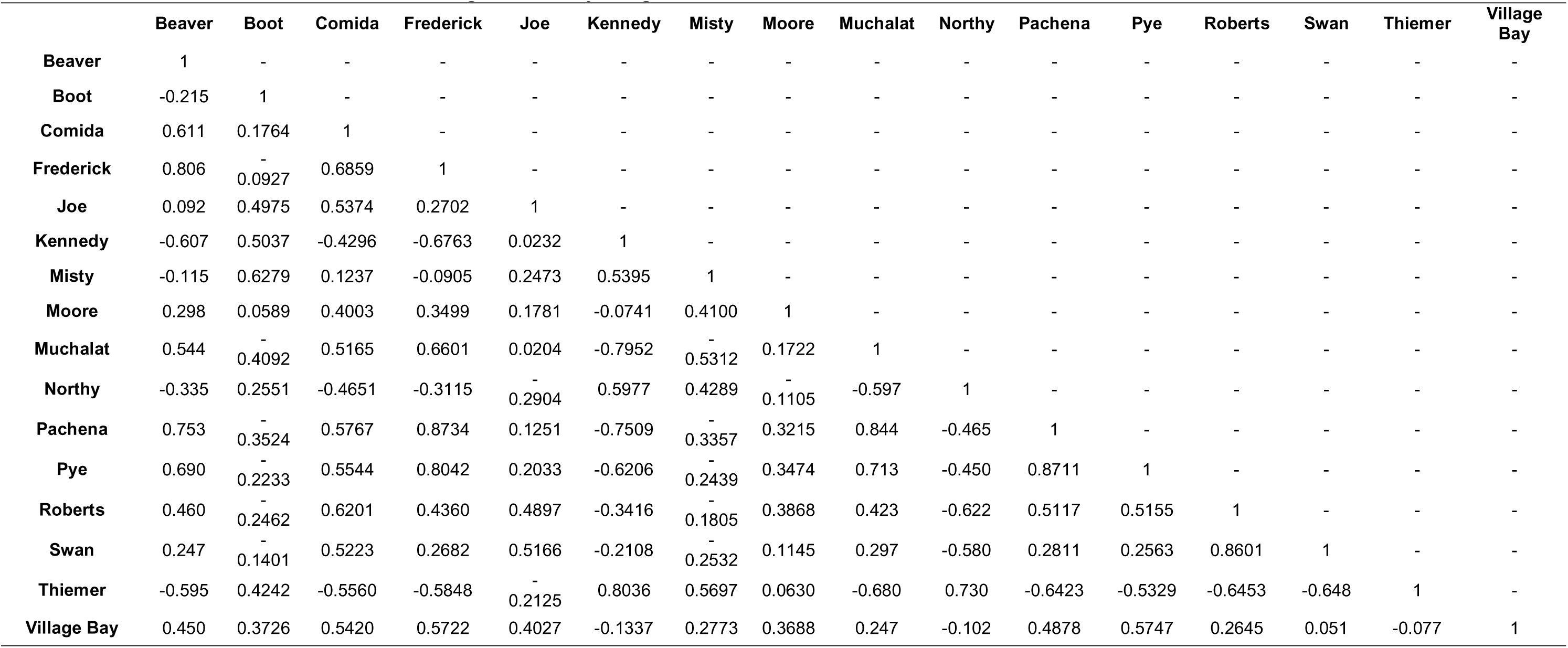
Matrix C of vector correlations between lineage evolutionary change vectors from Stuart et al 2017 data

**Table S2.** Spectral decomposition of C from Stuart et al. 2017

| Eigenvalues |  | Eigenvectors, Q |  |  |  |  |  |  |  |  |  |  |  |  |  |  |  |
| --- | --- | --- | --- | --- | --- | --- | --- | --- | --- | --- | --- | --- | --- | --- | --- | --- | --- |
| V | lineage | q1 | q2 | q3 | q4 | q5 | q6 | q7 | q8 | q9 | q10 | q11 | q12 | q13 | q14 | q15 | q16 |
| 7.405 | Beaver | 0.29 | 0.06 | 0.22 | 0.03 | 0.53 | -0.08 | -0.36 | -0.12 | 0.41 | -0.14 | 0.42 | 0.11 | 0.09 | 0.19 | 0.03 | 0.09 |
| 3.010 | Boot | -0.12 | 0.45 | -0.06 | -0.42 | -0.14 | -0.19 | -0.02 | 0.16 | 0.45 | -0.28 | -0.27 | -0.03 | 0.30 | -0.24 | 0.13 | 0.10 |
| 1.983 | Comida | 0.27 | 0.27 | -0.09 | -0.07 | 0.13 | -0.34 | 0.02 | 0.49 | -0.18 | 0.19 | 0.23 | -0.45 | -0.34 | -0.13 | -0.03 | 0.07 |
| 0.998 | Frederick | 0.31 | 0.13 | 0.24 | -0.11 | 0.24 | -0.01 | 0.29 | -0.11 | 0.05 | -0.09 | -0.40 | 0.19 | -0.44 | 0.03 | 0.29 | -0.44 |
| 0.598 | Joe | 0.11 | 0.36 | -0.37 | -0.27 | -0.11 | 0.08 | 0.41 | -0.48 | -0.04 | 0.02 | 0.43 | 0.04 | -0.04 | 0.13 | -0.12 | -0.03 |
| 0.543 | Kennedy | -0.29 | 0.22 | -0.14 | 0.12 | 0.05 | 0.37 | -0.10 | 0.31 | 0.37 | 0.18 | 0.11 | 0.16 | -0.23 | -0.08 | -0.38 | -0.42 |
| 0.347 | Misty | -0.14 | 0.46 | 0.09 | 0.19 | 0.16 | -0.32 | -0.07 | -0.11 | -0.20 | 0.56 | -0.20 | 0.31 | 0.26 | 0.11 | -0.03 | 0.04 |
| 0.292 | Moore | 0.12 | 0.30 | 0.13 | 0.70 | -0.29 | -0.20 | 0.13 | -0.08 | 0.08 | -0.43 | -0.02 | -0.12 | -0.01 | 0.04 | -0.22 | -0.04 |
| 0.199 | Muchalat | 0.31 | -0.15 | 0.13 | -0.06 | -0.34 | -0.10 | 0.27 | 0.46 | 0.09 | 0.11 | 0.23 | 0.30 | 0.35 | 0.32 | 0.08 | -0.24 |
| 0.179 | Northy | -0.26 | 0.11 | 0.29 | 0.03 | 0.47 | 0.30 | 0.52 | 0.19 | -0.18 | -0.16 | 0.06 | -0.21 | 0.33 | 0.02 | -0.05 | 0.06 |
| 0.138 | Pachena | 0.33 | -0.02 | 0.23 | 0.01 | -0.06 | 0.17 | 0.19 | 0.03 | 0.04 | 0.11 | 0.03 | 0.40 | -0.09 | -0.57 | -0.27 | 0.44 |
| 0.122 | Pye | 0.31 | 0.06 | 0.22 | -0.01 | -0.18 | 0.35 | -0.01 | -0.17 | 0.31 | 0.43 | -0.24 | -0.53 | 0.12 | 0.18 | -0.08 | 0.07 |
| 0.068 | Roberts | 0.27 | 0.09 | -0.36 | 0.33 | 0.09 | 0.27 | -0.09 | 0.00 | -0.07 | 0.06 | 0.09 | -0.04 | 0.32 | -0.42 | 0.49 | -0.23 |
| 0.053 | Swan | 0.21 | 0.03 | -0.53 | 0.13 | 0.20 | 0.20 | 0.08 | 0.25 | 0.05 | -0.11 | -0.35 | 0.18 | -0.07 | 0.41 | -0.04 | 0.42 |
| 0.039 | Thiemer | -0.31 | 0.18 | 0.20 | 0.15 | -0.23 | 0.18 | 0.07 | 0.08 | 0.15 | 0.11 | 0.24 | 0.08 | -0.33 | 0.13 | 0.60 | 0.35 |
| 0.027 | Village Bay | 0.16 | 0.38 | 0.22 | -0.21 | -0.18 | 0.40 | -0.44 | 0.10 | -0.50 | -0.25 | 0.01 | 0.10 | 0.01 | 0.15 | -0.05 | -0.03 |

**Table S3.**
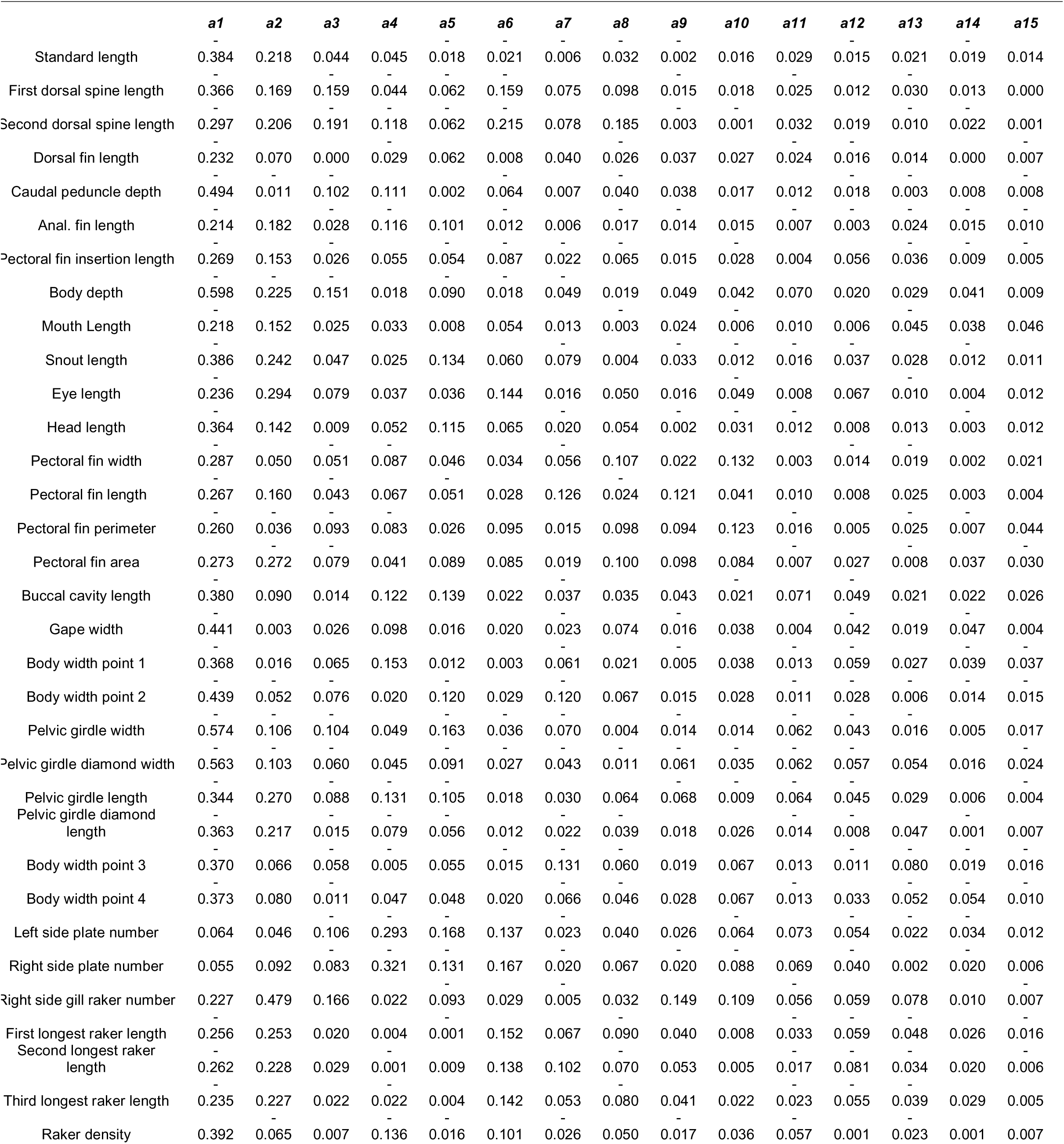
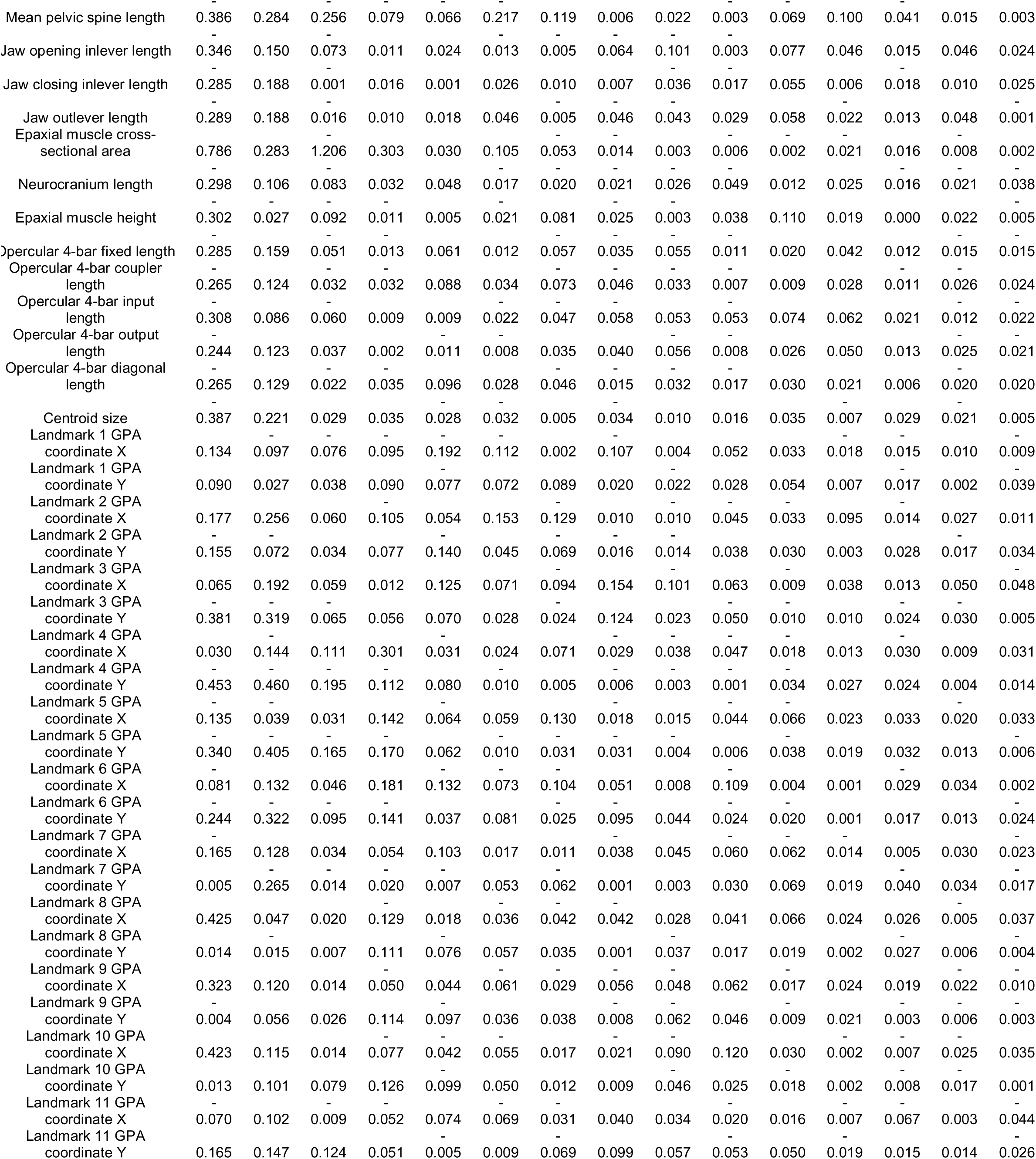
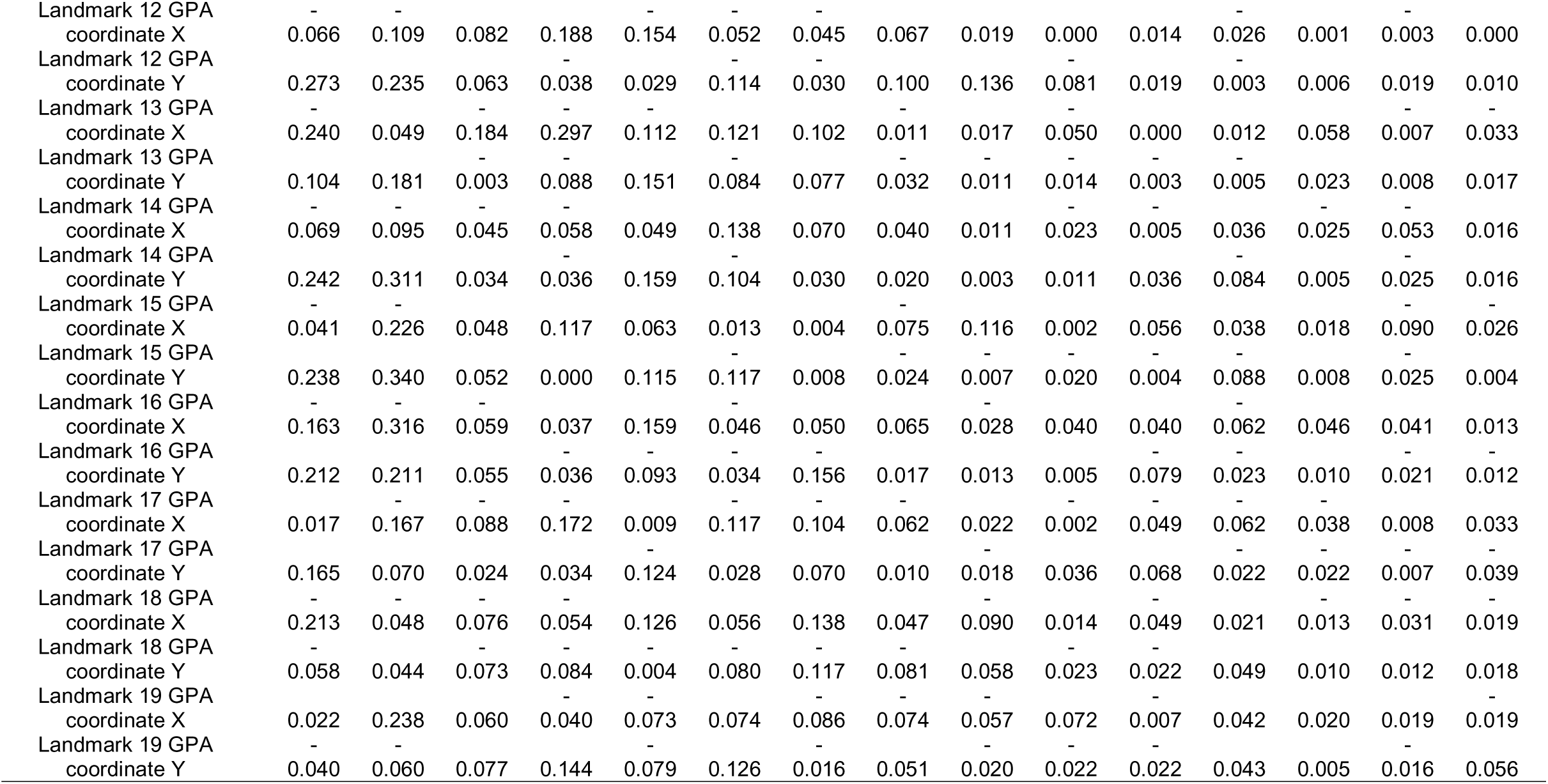
Matrix A relating eigenvectors of C to trait space

