## Supplemental SAS Script for "A Multivariate View of Parallel Evolution"

```
proc iml;
/* supplemental data table 1 from Stuart et al. (trait differences in units of phenotypic sd's)*/
xr = {-3.92	-9.73	-8.33	-2	-6.44	-1.06	-2.99	-9.12	-1.45	-0.61	-0.61	-1.72	-3.57	-4.63	-4.75	4.76	1.69	-6.35	-4.14	-8.64	-13.72	-12.08	-11	-5.06	-1.63	-2.98	-3.43	-3.09	5.48	-3.98	-3.99	-3.21	2.55	-10.5	-3.57	-3.01	-2.36	0.46	-3.08	-3.93	-1.78	-0.5	0.1	-4.05	-0.27	-4.07	-2.89	-1.64	6.57	-4.25	3.81	-5.09	-1.11	-8.12	-5.78	-5.96	-0.28	-3.03	1.35	1.3	9.41	-2.78	5.12	-4.3	2.92	-3.13	2.33	6.94	-2.9	9.8	-9.5	4.8	-2.6	8.47	4.14	7.46	-0.52	2.79	0.5	-1.3	-6.85	2.35	-0.07	-2.8,
6.77	10.3	14.37	4.15	1.29	7.2	5.37	-1.06	3.36	6.66	7.2	4.57	3.87	6.55	3.3	-9.6	1.57	0.86	-0.75	2.97	2.23	1.68	13.65	11.21	2	2.19	9.03	9.95	14.05	5.6	5.83	5.49	-1.99	15.69	1.48	4.27	2.64	-2	3.24	-0.82	4.22	3.85	5.25	3.69	4.25	7.61	-0.83	-4.7	5.91	-2.76	-0.45	-4.76	-12.04	-4.94	-0.62	-3.28	2.78	-3.04	4.14	-9.23	1.15	-5.09	0.22	-3.31	-1.56	-1.78	4.2	3.23	-4.32	10.91	11.9	5.77	-0.8	7.17	-6.3	7.88	-9.25	7.82	1.64	-2.13	-1.49	6.06	2.97	-4.38,
-1.62	1.91	4.81	-1.63	-3.78	0.74	-1.83	-6.52	-1.14	-1.21	-1.57	-1.87	0.71	-2.47	-1.2	0.62	-1.4	-2.07	-2.28	-3.15	-5.18	-5.73	0.5	-2.2	-1.45	-2.72	1.34	1.51	2.86	-2.14	-1.73	-1.89	2.67	0.51	-2.07	-0.47	-1.12	10.07	-1.56	-1.79	-1.53	-1.7	-3.32	-0.66	-1.16	-1.68	0.36	0.71	0.6	-3.22	3.96	-4.06	-2.71	-7.13	-4.07	-6.14	0.36	-6.81	-1.77	-1.92	3.8	-0.41	3.62	-0.43	6.1	0.05	2.49	3.14	-0.83	5.52	-3.22	3.29	-1.88	7.92	-4.05	6.93	-0.9	3.21	0.32	2.51	-0.97	-1.65	-0.38	-2.97,
-6.78	-7.7	-7.36	-1.8	-12.91	0.24	-4.47	-21.9	-0.58	-5.46	0.88	-5.05	-6.67	-3.32	-4.77	2.38	-8.96	-11.52	-10.53	-16.93	-17.06	-16.89	-7.93	-9.48	-9.28	-6.86	-2.48	-2.06	9.11	0.5	0.23	0.47	9.89	-3.23	-8.09	-4.65	-6.86	-4.31	-5.6	-9.79	-6.13	-3.98	-9.42	-4.68	-4.1	-7.53	-1.55	0.55	9.37	-8.17	0.98	-11.14	2.57	-13.78	-1.51	-11.53	-4.55	-7.87	1.35	-4.95	8.17	-1.68	9.53	-0.73	9.61	0.88	1.28	2.63	-6.55	9.34	-7.45	8.08	0.39	8.96	-3.46	9.44	-7.39	3.87	0.1	1.13	0.9	-0.88	1.64	-0.76,
1.18	2.51	3.65	0.8	-1.26	3.37	1.82	-3.66	1.11	-1.51	1.43	-2.39	-2.04	4.05	-1.5	-3.7	-2.87	-5.73	-6.14	-5.01	-5.18	-5.07	3.86	2.89	-0.86	-1.65	6.93	7.04	4.18	5.32	5.04	4.86	0.31	7.82	4.4	2.45	3.84	22.67	1.34	3.67	1.97	0.86	2.24	0.45	0.99	1.9	4.85	2.23	2.53	-3.02	-2.33	-9.84	-5.48	-5.7	-1.85	-4.12	-3.46	-1.67	-0.42	-2.81	4.8	0.04	3.75	0.47	5.64	-0.73	0.62	-1.65	-6.81	1.41	8.65	5.17	-0.2	5.09	-3.84	4.81	-5.62	-0.53	-1.15	4.18	0.37	1.42	4.13	-5.14,
8.92	7.1	7.73	5.7	9.43	6.1	5.28	10.64	7.03	10.58	12.71	11.02	10.8	10.78	11.5	-0.06	10.41	10.29	9.04	10.44	8.26	6.08	6.97	6.56	9.28	10.61	2.03	1.98	5.69	10.47	9.78	9.19	-5.67	8.14	5.34	7.61	7.22	4.43	8.49	5.1	7.12	6.14	7.55	5.46	6.54	8.95	-7.6	1.67	3.87	2.51	5.31	7.11	-3.32	1.88	-0.07	-0.22	2.48	0.96	4.76	-0.67	-9.34	0.56	-6.12	0.97	-9.69	1.89	4.9	2.36	0.79	-0.58	5.61	-3.99	3.69	-3.75	1.08	-4.76	-1.65	-1.9	-6.66	-0.42	4.89	0.33	3.37	2.57,
6.31	5.31	4.45	2.78	3.39	4.13	4.74	-0.82	2.72	6.45	4.39	3.98	3.21	3.69	0.12	-5.27	3.23	2.95	3.79	0.32	0.86	1.19	4.47	5.48	4.02	4.82	-7.61	-6.83	9.59	3.17	2.38	2.92	-4.29	8.17	5.57	3.76	4.92	-1.99	2.96	1.2	4.89	3.68	2.38	3.86	3.4	6.36	-2.13	-0.51	2.12	-0.1	5.62	-4.98	-1.33	-5.48	-0.23	-5.13	1.7	-6.17	2.91	-5.04	-0.73	-0.42	3.41	2.44	2.43	2.36	1.38	0.15	-0.97	0.96	-3.34	3.71	-3.95	4.82	-3.05	5.47	-2.74	2.89	-3.39	-0.67	0.48	1.62	5.64	-1.39,
0.94	-0.03	-0.43	-0.67	0.31	-1.33	1.44	-2.22	0.48	-0.09	1.08	-0.09	-2.81	1.21	-3.11	-1.61	0.47	-0.19	0.95	-0.95	-0.59	-0.35	-0.51	-0.53	-1.31	-0.65	-2.37	-2.63	2.92	0.79	0.7	0.21	-1.68	0.35	-0.23	0.43	0.05	3.18	0.34	-1.06	-0.46	-1.07	0.16	0.25	-0.96	0.94	-0.34	1.52	-0.37	0.25	-0.33	-2.02	3.31	-5.04	1.72	-5.06	3.86	-3.76	1.12	-1.93	-0.83	1.26	0.15	1.55	-0.14	2.11	-0.11	2.1	2.35	2.08	-3.07	-0.13	-1.85	1.21	0.08	2.28	-2.29	1.37	-2.55	1.55	-2.05	-0.74	0.32	1.11,
-4.17	-1.9	0.42	-2.85	-5.06	-4.1	-3.57	-2.41	-2.59	-6.69	-2.58	-4.6	-2.02	-1.19	-1.01	4.38	-4.59	-3.81	-4.17	-3.44	-4.61	-4.71	-3.15	-4.21	-7.06	-6.12	1.59	1.57	3.34	-5.23	-4.61	-4.58	3.89	-2.99	-3.78	-3.02	-3.67	-0.67	-4.47	-1.5	-4.66	-5.24	-3.54	-1.42	-4.32	-3.93	2.53	2.36	-0.31	0.11	0.34	-2.2	0.79	-2.95	-0.37	-2.42	-0.45	-2.71	-3.8	1.17	3.9	0.62	1.59	0.66	3.93	0.49	-1.84	2.44	2.05	3.36	-0.68	-0.66	-0.63	-0.62	-0.8	-1.43	0.19	0.38	-1.47	2.98	-2.09	-1.79	-0.73	1.14,
3.09	0.83	0.39	2.81	1.8	2.54	-0.43	0.25	2.24	5.26	4.7	4.97	1.42	2.49	1.31	-1.58	4.92	3.89	2.95	-0.36	-0.45	2.18	2.67	2.07	-0.22	1.23	-2.06	-0.4	-0.55	3.82	4.65	3.45	-2.69	-0.62	2.44	2.9	3.03	-12.74	1.79	2.04	2.05	1.89	0.82	2.91	2.78	2.93	-5.73	0.69	-0.38	-1.11	2.39	2.36	1.05	1.07	2.64	-0.02	-1.71	0.19	3.21	-1.82	-3.58	-0.07	-2.41	-0.03	-3.4	0.26	0.25	0.16	-1.57	-1.36	1.61	2.1	2.23	1.64	0.12	0.92	3.09	-3.78	-4.08	-0.98	4.73	-3.19	-1	0.72,
-12.1	-14.35	-13.25	-7.4	-18.63	-7.27	-7.32	-22.73	-7.56	-13.82	-2.37	-11.24	-10.22	-7.01	-7.29	10.09	-15.76	-11.99	-10.48	-14.21	-22.65	-22.77	-12.35	-9.52	-11.5	-6.41	0.99	1.2	13.65	-7.19	-7.59	-7.16	13.21	-15.72	-14.54	-9.34	-11.81	-2.36	-10.57	-8.93	-8.75	-8.17	-8.27	-2.36	-8.69	-11.47	0.97	8.09	6.7	-3.96	3.19	-10.7	3.97	-17.49	2.28	-14.22	-5.52	-8.89	-6.62	0.75	13.06	3.45	9.19	3.38	13.77	4.09	-3.77	5	-1.69	4.65	-6.27	0.91	4.49	4.72	7.4	3.92	-11.14	5.33	-8.57	8.62	-4.26	-7.07	1.43	2.94,
-6.91	-14.88	-18.27	-2.48	-12.46	-6.05	-9.98	-15.73	-2.85	-6.69	-4.34	-9.27	-3.31	-0.95	-1.2	8.31	-8.63	-8.55	-4.26	-7.58	-11.57	-12.27	-5.43	-3.55	-7.45	-7.28	4.12	2.68	18.29	4.83	4.66	4.9	12.85	-19.44	-13.5	-8.05	-4.01	3.08	-12.39	-10.15	-10.37	-8.53	-11.68	-9.14	-9.21	-7.38	4.26	4.66	4.97	-0.13	1.26	-15.06	0.09	-17.36	-1.01	-12.42	-4.54	-4.67	-1.75	-0.47	9.74	2.04	6.83	2.11	10.66	4.1	-4.86	5.46	0.21	4.2	-4.39	-0.88	-3.06	5.26	-1.4	6.11	-7.18	9.81	-0.82	7.03	-9.52	-4.17	6.41	0.71,
-4.64	-8.75	-6.72	-4.46	-6.44	-4.99	-5.69	-9.54	-3	-2.52	-1.19	-2.51	-3.87	-1.35	-1.59	6.06	-1.77	-3.87	-3.65	-6.69	-9.6	-10.1	-6.75	-7.2	-6.93	-4.77	0.45	-0.11	4.11	-4.09	-4.17	-3.62	5.77	-8.76	-3.14	-2.98	-3.44	60.83	-2.9	-5.41	-2.25	-1.34	-4.38	-5.12	-1.53	-5.12	-2.13	2.22	4.18	1.01	1.4	-5.4	3.9	-8.65	0.62	-9.03	-1.75	-7.27	-3.78	-2.74	7.08	2.19	6.71	2.68	8.71	2.98	-5.67	1.86	-1.02	1.81	-5.86	0.82	1.53	5.46	1.74	4.93	-1.84	2.86	-4.21	2.35	-2.02	-1.65	1.13	4.95,
-7.87	1	0.26	-2.34	-3.5	-1.97	-3.37	-4.55	-1.06	-4.06	-0.88	-0.98	-1.4	-1.97	1.21	10.23	-2.94	-6.07	-6.84	-3.36	-5.3	-5.78	-0.38	-3.75	-3.46	-3.96	-0.59	0.35	-0.05	-1.61	-2.32	-1.81	5.58	3.24	-2.77	-3.25	-2.92	48.41	-1.46	-2.31	-1.91	-2.08	-3.06	-3.67	-2.47	-7.63	-3	0.18	3.93	-5.81	4.97	0.21	-0.35	0.51	1.01	0.21	-0.84	-0.75	0.89	0.07	1.48	0.82	0.79	-0.77	0.6	-2.01	-0.4	-0.67	-4.36	2.19	1.28	1.94	2.74	1.5	-3.91	0.69	0.84	0.12	-0.79	-0.95	1.85	-0.69	-3.24	-0.51,
8.17	5.68	3.47	2.69	6.62	2.11	5.8	7.75	8.13	6.39	8.8	6.36	5.14	7.01	5.57	-5.78	7.55	7.07	4.84	6.89	7.95	4.52	5.96	7.72	6.04	6.88	0.16	-0.05	3.9	7.63	7.7	7.5	-6.29	4.96	6.27	7.64	6.54	-18.36	2.29	5.14	3.46	4.22	4.57	4.68	4.02	7.99	-3.08	1.58	-0.11	4.79	-0.33	2.73	2.96	-1.21	4.96	-3.21	5.31	-2.19	1.93	-0.68	-5.29	2.76	-6.74	2.82	-4.46	3.45	-0.07	0.13	3.47	-2.04	0.76	-2.76	1.23	-2.44	-0.75	-2.77	-2.38	-2.18	-4.77	0.4	-0.83	-1.14	1.69	4.86,
0.26	-3.3	-2.78	-2.15	-6.69	0.88	-0.77	-7.64	0.96	2.93	5.05	0.46	-0.99	-4.03	-3.74	-4.48	-2.36	-2.73	-2.13	-1.39	-4.34	-3.29	3.74	1.35	-1.53	-2	5.66	8	6.77	1.98	0.65	1.91	4.58	-4.35	1.07	1.79	1.79	-1.27	-0.53	-4.57	1.89	1.57	-1.26	1.2	1.39	-0.22	-1.69	0.97	10.67	0.19	7.28	-6.81	-2.69	-10.17	-3.01	-7.49	2.99	-4.96	-0.23	-1.78	3.26	0.38	3.42	1.32	4.17	2.83	0.09	4.71	-1.82	2.96	1.07	0.47	-1.13	2.61	-6.68	3.62	-8.06	6.96	-2.5	1.93	-2.69	0.17	7.43	1.75};
X = J(16, 84);
do i = 1 to 16;
vt = xr[i,];
X[i,] = vt/sqrt(sum(vt#vt)); /* normalize row vectors*/
end;
cor = X*X`; /* equation 5 */
vec = eigvec(cor);
A = X`*inv(vec`); /* equation 7 */
A1 = A[,1]/sqrt(sum(A[,1]#A[,1])); /* I normalized these A vectors, but same results with raw values */
A2 = A[,2]/sqrt(sum(A[,2]#A[,2]));
A3 = A[,3]/sqrt(sum(A[,3]#A[,3]));
ob = eigval(cor);
print A;
print ob;
print vec;
print cor;
create observed from ob;
append from ob;
close observed;
create popvec from vec[colname={"PC1" "PC2" "PC3" "PC4" "PC5" "PC6" "PC7" "PC8" "PC9" "PC10" "PC11" "PC12" "PC13" "PC14" "PC15" "PC16"}];
append from vec;
close popvec;
t = nrow(a1);
A13 = A1||A2||A3;
create Amat from A13[colname={"A1" "A2" "A3"}];
append from A13;
close Amat;
run;
data Amat;
set  Amat;
rank=_n_;


 /* Construct the distribution of eigenvalues under the null, using the Wishart distribution */
%let N = 84;              /* sample size NUMBER OF TRAITS */
%let NumSamples = 1000;
proc iml;
start Cov2Corr(A);    /*this is just a function to convert a cov to corr matrix*/
   D = sqrt(vecdiag(A));
   return( A / D` / D );
finish;
proc iml;
NumSamples = &NumSamples;
DF = &N - 1;              /* X ~ N obs from MVN(0, Sigma). In this case N is number of traits */
Sigma = I(16);  /* null hypothesis */
S = RandWishart(NumSamples, DF, Sigma); /* each row is 16x16 matrix */
X = J(nrow(S), 16);
do i = 1 to nrow(S);
A = shape(S[i,], 16, 16);
D = sqrt(vecdiag(A));
CorrS = ( A / D` / D );
D = eigval(CorrS);
X[i,] = D`;
end;
create results2 from X[colname={"Eigen1" "Eigen2" "Eigen3" "Eigen4" "Eigen5" "Eigen6" "Eigen7" "Eigen8" "Eigen9" "Eigen10" "Eigen11" "Eigen12" "Eigen13" "Eigen14" "Eigen15" "Eigen16"}];
append from X;
close results2;
proc transpose data = results2 out = tresults2;
proc sort data = tresults2;
by _NAME_;
proc transpose data = tresults2 out = long2;
by _NAME_;
data  long2;
set  long2;
if _NAME_  = "Eigen1" then rank = 1;
if _NAME_  = "Eigen2" then rank = 2;
if _NAME_  = "Eigen3" then rank = 3;
if _NAME_  = "Eigen4" then rank = 4;
if _NAME_  = "Eigen5" then rank = 5;
if _NAME_  = "Eigen6" then rank = 6;
if _NAME_  = "Eigen7" then rank = 7;
if _NAME_  = "Eigen8" then rank = 8;
if _NAME_  = "Eigen9" then rank = 9;
if _NAME_  = "Eigen10" then rank =10;
if _NAME_  = "Eigen11" then rank = 11;
if _NAME_  = "Eigen12" then rank = 12;
if _NAME_  = "Eigen13" then rank = 13;
if _NAME_  = "Eigen14" then rank = 14;
if _NAME_  = "Eigen15" then rank = 15;
if _NAME_  = "Eigen16" then rank = 16;
proc sort data = long2;
by rank;

 /* Construct the distribution of eigenvalues under the null, using random vectors in trait space */
proc iml;
vals = J(1000, 16);
do k = 1 to 1000;
X = J(16, 84);
do j = 1 to 16;
z = j(84,1);
call randgen(z, "Normal"); /* random vector */
v = z`;
vs = v/sqrt(sum(v#v));
X[j,] = vs;
A = X*X`;
D = sqrt(vecdiag(A));
xcor = ( A / D` / D );
ve = vech(xcor);
end;
val = eigval(xcor);
vals[k,] = val`;
end;
create results from vals[colname={"Eigen1" "Eigen2" "Eigen3" "Eigen4" "Eigen5" "Eigen6" "Eigen7" "Eigen8" "Eigen9" "Eigen10" "Eigen11" "Eigen12" "Eigen13" "Eigen14" "Eigen15" "Eigen16"}];
append from vals;
close results;

/* some ugly code to merge the observed and two null datasets */
proc transpose data = results out = tresults;
proc sort data = tresults;
by _NAME_;
proc transpose data = tresults out = long;
by _NAME_;
data  long;
set  long;
if _NAME_  = "Eigen1" then rank = 1;
if _NAME_  = "Eigen2" then rank = 2;
if _NAME_  = "Eigen3" then rank = 3;
if _NAME_  = "Eigen4" then rank = 4;
if _NAME_  = "Eigen5" then rank = 5;
if _NAME_  = "Eigen6" then rank = 6;
if _NAME_  = "Eigen7" then rank = 7;
if _NAME_  = "Eigen8" then rank = 8;
if _NAME_  = "Eigen9" then rank = 9;
if _NAME_  = "Eigen10" then rank =10;
if _NAME_  = "Eigen11" then rank = 11;
if _NAME_  = "Eigen12" then rank = 12;
if _NAME_  = "Eigen13" then rank = 13;
if _NAME_  = "Eigen14" then rank = 14;
if _NAME_  = "Eigen15" then rank = 15;
if _NAME_  = "Eigen16" then rank = 16;
group = "emprical null";
proc sort data = long;
by rank;
data observed;
set  observed;
rank=_n_;
data observed(rename=(COL1=observe));
do repno=1 to 1000;
  do _n_=1 to n;
    set observed nobs=n point=_n_;
    output;
  end;
end;
stop;
proc sort data = observed;
by rank;
data long2(rename=(COL1=theor));
group = "theoretical null(Wishart)";
set long2;
data total;
merge long long2 observed;


/* figure 4 */
proc sgplot data=total noautolegend;
vbox COL1/ DISCRETEOFFSET = .2 nooutliers nocaps  CONNECTATTRS=(Color="green" Thickness=2) category = rank  fill fillattrs = (color = green) legendlabel="Null distribution (empirical)";
vbox theor/DISCRETEOFFSET = -.2 nooutliers nocaps  CONNECTATTRS=(Color="yellow" Thickness=2) category = rank  fill fillattrs = (color = yellow) legendlabel="Null distribution (Wishart)";
vbox observe/nocaps connect = mean CONNECTATTRS=(Color="blue" Thickness=2) category=rank fill fillattrs = (color = blue) legendlabel="Observed distribution";
keylegend / location=inside noborder position=topright;
xaxis label = "Eigenvalue rank" labelattrs =(size = 14 weight = bold);
yaxis  label = "Eigenvalue" labelattrs =(size = 14pt weight = bold);

/* add in trait and population info */
data traits;
input type $	rank;
datalines;
S	1
D	2
D	3
S	4
S	5
S	6
S	7
S	8
U	9
U	10
U	11
U	12
S	13
S	14
S	15
S	16
T	17
T	18
S	19
S	20
D	21
D	22
D	23
D	24
S	25
S	26
D	27
D	28
T	29
T	30
T	31
T	32
T	33
D	34
T	35
T	36
T	37
T	38
T	39
T	40
T	41
T	42
T	43
T	44
T	45
U	46
U	47
U	48
U	49
U	50
U	51
U	52
U	53
U	54
U	55
U	56
U	57
U	58
U	59
U	60
U	61
U	62
U	63
U	64
U	65
U	66
U	67
U	68
U	69
U	70
U	71
U	72
U	73
U	74
U	75
U	76
U	77
U	78
U	79
U	80
U	81
U	82
U	83
U	84
;
data traitaxis;
merge Amat traits;
data traitaxis;
set traitaxis;
aA1 = abs(A1);
aA2 = abs(A2);
aA3 = abs(A3);
if type = "S" then type2 = "Swimming";
if type = "D" then type2 = "Defense";
if type = "T" then type2 = "Trophic";
if type = "U" then type2 = "Unclassified";
data popvec;
set  popvec;
rank=_n_;
data popvec;
set popvec;
if rank = 1 then pop = "Beaver";
if rank = 2 then pop = "Boot";
if rank = 3 then pop = "Comida";
if rank = 4 then pop = "Frederick";
if rank = 5 then pop = "Joe";
if rank = 6 then pop = "Kennedy";
if rank = 7 then pop = "Misty";
if rank = 8 then pop = "Moore";
if rank = 9 then pop = "Muchalat";
if rank = 10 then pop = "Northy";
if rank = 11 then pop = "Pachena";
if rank = 12 then pop = "Pye";
if rank = 13 then pop = "Roberts";
if rank = 14 then pop = "Swan";
if rank = 15 then pop = "Thiemer";
if rank = 16 then pop = "Village";

/* figure 6 */
proc sgplot data=traitaxis noautolegend;
vbox A1/ DISCRETEOFFSET = .1 nooutliers boxwidth = .1 category = type2  fill fillattrs = (color = green) transparancy = .4 legendlabel="Dimension 1";
vbox A2/DISCRETEOFFSET = 0  nooutliers  boxwidth = .1 category = type2 fill fillattrs = (color = yellow) transparancy = .1 legendlabel="Dimension 2";
vbox A3/DISCRETEOFFSET = -.1  nooutliers  boxwidth = .1 category = type2 category=rank fill fillattrs = (color = blue) transparancy = .4 legendlabel="Dimension 3";
keylegend / location=inside noborder position=topleft;
xaxis label = "Trait type" labelattrs =(size = 14 weight = bold);
yaxis  label = "Loading on vector of parallel evolution" labelattrs =(size = 14pt weight = bold);

/* figure 5 */
proc sort data = popvec;
by PC1;
proc sgplot data=popvec noautolegend;
vbar pop/ response = PC1 CATEGORYORDER=RespDesc;
xaxis discreteorder=data  label = "Population" labelattrs =(size = 14 weight = bold);
yaxis  label = "Loading on PC1 of C" labelattrs =(size = 14pt weight = bold);
proc sort data = popvec;
by PC2;
proc sgplot data=popvec noautolegend;
vbar pop/ response = PC2 CATEGORYORDER=RespDesc;
xaxis discreteorder=data  label = "Population" labelattrs =(size = 14 weight = bold);
yaxis  label = "Loading on PC2 of C" labelattrs =(size = 14pt weight = bold);
proc sort data = popvec;
by PC3;
proc sgplot data=popvec noautolegend;
vbar pop/ response = PC3 CATEGORYORDER=RespDesc;
xaxis discreteorder=data  label = "Population" labelattrs =(size = 14 weight = bold);
yaxis  label = "Loading on PC3 of C" labelattrs =(size = 14pt weight = bold);
run;
```
